# Gregariousness can promote invasion success of pest insects while higher environmental heterogeneity decreases it

**DOI:** 10.64898/2026.11.05.764464

**Authors:** M. Jonsson, V. Korhonen, E. Guilbault, K. Koskenpato, S. Van Meyel, C. Lindstedt

## Abstract

Climate change and environmental homogenization are expected to facilitate the expansion of many forest pest insect species and increase the vulnerability of forest ecosystems to insect outbreaks. At present, we lack empirical data on how the antipredator defence traits of pest insects interact with the local environmental conditions and shape outbreak risk. Here we study this and ask 1) how the antipredator defence strategy of pest insects shapes survival under naturally varying predator community, and 2) how is predation risk shaped by the diversity and structure of forests? We tested these questions by conducting a field predation experiment with artificial prey system (prey items made from edible dough) where we manipulated the gregariousness, unpalatability and defensive colouration of prey and tested their effects on prey survival. Additionally, we measured environmental variables and invertebrate predator community structure along the prey transects to analyse how they shape overall predation risk. Our results show that the predation risk was always lowest for gregarious prey irrespective of other defence strategies. Furthermore, our results indicate that predation pressure was lower in more homogenously structured forest areas or areas with a weak quality deadwood continuum in comparison to locations with more natural state forests and good quality deadwood continuum. Altogether, our results suggest that gregariousness can facilitate pest insects to form outbreaks by increasing their survival against generalist predators. However, by maintaining higher heterogeneity in forests, it is possible to ensure higher predation pressure to regulate their population growth.

## Introduction

Global changes such as climate change, biodiversity loss, habitat homogenisation, and invasive species are rapidly altering the spatial distribution ranges of organisms and disrupting species interactions (Biedermann et al., 2019; Jaime et al., 2024). For example, such dynamics often place species and populations in new environments (e.g., novel competing species or predators) to which they have not previously adapted (HilleRisLambers et al., 2013). This can lead to species extinctions and the introduction of novel pathogens and pests, resulting in cascading impacts on e.g., global food security (Subedi et al., 2023), the availability of natural resources (Morris et al., 2017), and the health of both human and non-human organisms (Wagner et al., 2021). A key challenge is to identify the factors that enable ecosystems to resist and recover from these rapid changes while maintaining their ecological processes. This requires information about the characteristics that underlie the success of invading organisms in the habitats they invade and how these are shaped by ecological conditions, such as the local predator community.

One important characteristic that has been suggested to facilitate invasion of novel environments is sociality, such as cooperation (Devegili et al., 2025; Komdeur & Ma, 2021) or group living (Wrensford et al., 2025). For example, cooperative breeding has been shown to enhance the invasion success of birds in harsh environments (Cornwallis et al., 2017), and the transmission of novel information via social learning has been demonstrated to impact how individuals learn about the quality of new food sources (Thorogood et al., 2018). Many agricultural pest insects (i.e., species attacking or infesting crops or plantations) also benefit from sociality. For example, desert locusts (*Schistocerca gregaria*) switch from a solitary to gregariously feeding stage as a response to increased predation risk at higher population densities (Sword et al., 2000). However, the significance of social traits in the success of pest insects in forests remains largely unknown (Biedermann & Taborsky, 2011; Smith et al., 2018). This is despite the fact that many forest pests have evolved a variety of social behaviours, such as cooperation within groups in antipredator defence (Ritter et al., 2026; Van Meyel et al., 2026), division of labour in colonies (Biedermann, 2020), thermoregulation via aggregations (Seymour, 1974) and group facilitated feeding (Ghent 1960; Peterson 1987).

Another partly neglected factor in predicting the success of forest pest insects is their antipredator defence strategies and how these can shape their survival against predators (Hunter, 1991; Lindstedt et al., 2022). For instance, aposematic species (including insects) often possess toxic and/or unpalatable chemical compounds that they advertise with warning colours to potential predators (Poulton, 1890; Cott, 1940; Mappes et al., 2005, Ruxton et al., 2018). Predators can learn to associate the unprofitability of such prey with the conspicuous and easily memorable signals and avoid attacking them in future encounters (Sherratt & Beatty, 2003; Stevens & Ruxton, 2012). This avoidance learning effect should be even stronger if the prey is group-living, as this increases the frequency with which predators encounter warning colours and enhances their association with chemical defences (Riipi et al., 2001; Sillén-Tullberg & Leimar, 1988). Additionally, group living may provide survival benefits to prey through the dilution effect; that is, a decreased per capita attack rate (Lehtonen & Jaatinen, 2016; Turner & Pitcher, 1986, Treisman, 1975).

Chemical defences, combined with different types of protective colouration strategies and group living, are also common among forest pest insects. For example, many pine sawfly species defend themselves chemically against predator attacks by collectively deploying defensive fluids from their mouths, in conjunction with a defensive display (Lindstedt et al., 2025). In some of these species, such as the red- headed pine sawfly (*Neodiprion lecontei*), this display is also combined with conspicuous aposematic colouration which predators learn to avoid (Lindstedt et al., 2022; Linnen et al., 2018). However, majority of the population dynamic models of forest pest insects consider situations where the focal pests are palatable and live solitarily. Thus, predation risk is expected to increase with the density of prey individuals until a certain maximum level in the predators’ ability to consume the prey is reached (Holling, 1965; Delong, 2021). Yet, contrasting patterns may emerge for group-living, chemically defended, or aposematic species whose defence strategies are based on ‘strength in numbers’ (Ruxton & Sherratt, 2006).

To determine the mechanisms that influence on the survival of pest species in environments against natural predator community, it is also crucial to understand the ecosystem characteristics that make them more vulnerable to outbreaks. Previous research indicates that insect outbreaks are typically less intense in heterogeneous environments (Jactel & Brockerhoff, 2007; Jactel et al., 2021), where there is e.g., reduced availability of suitable host plants (Guo et al., 2019). Additionally, such environments—characterized by variations in tree age, species composition, and deadwood—often support higher densities and diversities of predators and parasitoids that regulate insect populations at various life stages (Blomqvist et al. 2016; Rappa et al., 2024). For instance, structural complexity in tree stands provides habitats for a broader range of species compared to homogeneous forests or monocultures (Huuskonen et al., 2021). Deadwood serves as nesting sites and shelter for various organisms, including birds (Vatka et al., 2014), ants (Lindgren, 2002), beetles (Gunnarsson et al., 2004), spiders (Varady-Szabo & Buddle, 2006), as well as bats and voles (Radu, 2006). The variation in vegetation can therefore foster a more diverse and dense predator community (Root, 1973; Russell, 1989), which can influence the efficacy of different antipredator strategies (Kikuchi et al., 2021; Medina et al., 2025). For example, predators can differ in their susceptibility to the chemical defences of prey (Burdfield-Steel et al., 2020; Exnerová et al., 2015), the amount of toxins predators have consumed already (Halpin et al., 2017; Speed & Ruxton 2014) as well as sensory and cognitive capabilities (Endler & Mappes, 2004; Marples & Kelly, 2000). Consequently, the efficacy of antipredator defence strategies employed by prey species may vary locally based on the abundance and composition of the predator community (see also Endler & Mappes, 2004; Medina et al., 2025). Given this variability among predators in their responses to defence strategies of prey, it is essential to evaluate how different strategies perform across the entire community to accurately identify the factors shaping the success of pest species in new environments.

In this study, our first goal was to examine how gregariousness, combined with chemical defence and aposematism, affects the survival of prey against the natural predator community (i.e., including both vertebrate and invertebrate predators). Secondly, we investigated how survival varied depending on vegetation structure, levels of forest management, and the predator community. We conducted a field predation experiment in boreal forests where we used artificial, caterpillar-mimicking, dough-made edible models as prey. We established eight treatment groups in which we manipulated gregariousness (solitary vs. groups of five prey), colouration (camouflaged = green prey against a green background vs. aposematic = black prey with yellow dots on the back against a green background), and palatability (unpalatable vs. palatable), in all possible combinations (see **Table 1**).

**Table 1.**
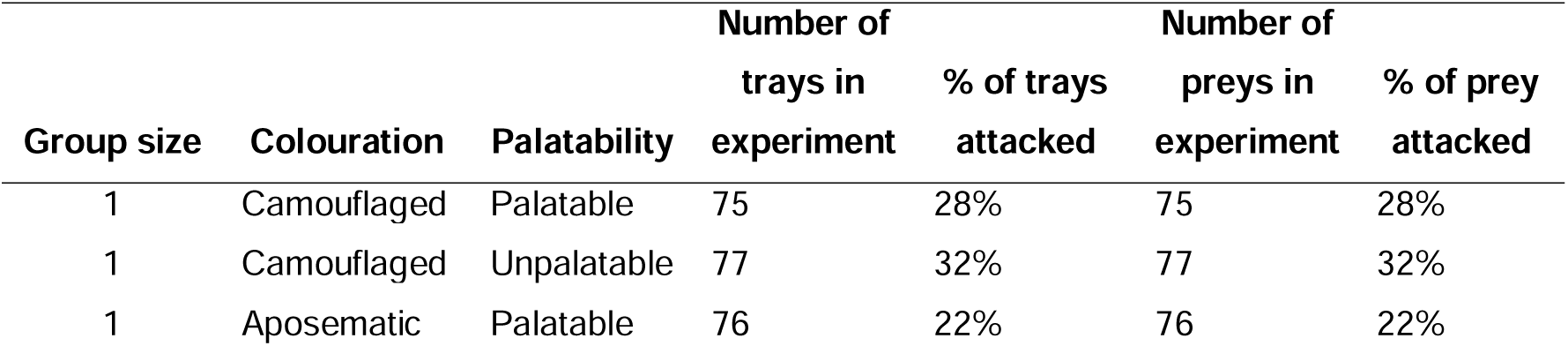
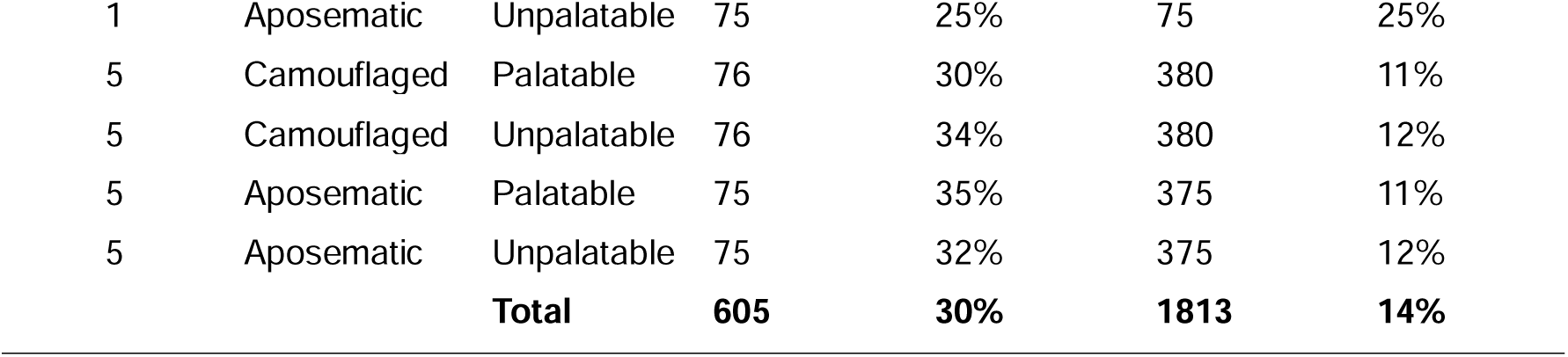
Final sample sizes for the eight different antipredator defence treatments groups and their respective initial attack rates during the experiment.

| **Group size** | **Colouration** | **Palatability** | **Number of trays in experiment** | **% of trays attacked** | **Number of preys in experiment** | **% of prey attacked** |
|---|---|---|---|---|---|---|
| 1 | Camouflaged | Palatable | 75 | 28% | 75 | 28% |
| 1 | Camouflaged | Unpalatable | 77 | 32% | 77 | 32% |
| 1 | Aposematic | Palatable | 76 | 22% | 76 | 22% |
| 1 | Aposematic | Unpalatable | 75 | 25% | 75 | 25% |
| 5 | Camouflaged | Palatable | 76 | 30% | 380 | 11% |
| 5 | Camouflaged | Unpalatable | 76 | 34% | 380 | 12% |
| 5 | Aposematic | Palatable | 75 | 35% | 375 | 11% |
| 5 | Aposematic | Unpalatable | 75 | 32% | 375 | 12% |
|  |  | **Total** | **605** | **30%** | **1813** | **14%** |

We tested six main hypotheses. Firstly (**I**), we expected that aposematically coloured prey would experience a lower attack risk compared to camouflaged green prey due to, for example, the neophobic responses of predators (Lindström et al., 1999; Marples & Kelly 2000) and/or their previous experience with similarly coloured but unpalatable natural prey (i.e., experienced predators have already learned to avoid warningly coloured prey, Lindström et al., 2001). This expectation arises from the fact that the predation experiment was conducted in the field, and we did not expect predators to be naïve to the aposematic prey types. Alternatively, if the predator community is dominated by naïve predators and/or those less sensitive to visual warning signals, the attack risk of aposematically coloured prey could be higher compared to camouflaged prey due to its increased detectability (Mappes et al., 2014; Koskenpato et al. 2026).

Secondly (**II**), we predicted that unpalatable prey would have a lower attack risk than palatable prey (Alatalo & Mappes, 1996), with the predation risk being even lower when unpalatability is combined with aposematic colouration (Lindström et al., 2001). Thirdly (**III**), we expected predation risk to be lower in groups of prey compared to solitary individuals. This is because the group itself can act as a warning signal (Mappes & Alatalo, 1996), increasing the reluctance of predators to attack groups. Lower predation risk can also result from the dilution effect (Riipi et al., 2001). Alternatively, larger groups may be easier to detect, potentially increasing the attack risk (Koskenpato et al., 2026; Lindström et al., 1999). Fourthly (**IV**), we predicted that being in a group would increase the attack risk for palatable camouflaged prey due to increased detectability, but would decrease the attack risk for aposematically coloured and unpalatable prey due to, for example, enhanced learning and/or signal stimulus (Riipi et al., 2001; McLellan et al. 2022).

In terms of environmental variables (**V**), we expected that predation risk across all antipredator defence treatments would increase with rising forest heterogeneity and with a higher quantity and quality of the deadwood, as these factors have shown to be associated with greater predator density and diversity (Bellone et al., 2017; Wrethling et al., 2026). Lastly (**VI**), we expected to observe fewer differences between the different antipredator defence treatments in more heterogeneous environments compared to more homogeneous environments. That is, environments with a more diverse and dense predator community are more likely to include predators that target defended prey (Klapwijk et al., 2026; Medina et al., 2025; Nokelainen et al., 2014). Alternatively, the predation risk can also be high in a more homogeneous environment if predators happen to be very effective at foraging (e.g., certain abundant generalist wood ant species) (see Lindstedt et al., 2006).

## Methods

### Field predation experiment

To estimate the predation risk of prey items with various antipredator defence strategies, we used artificial dough-made prey mimicking Lepidopteran caterpillars in shape (**Fig. S1**). Prey items were created using protocols adapted from J. B. Barnett *et al*. (2016) (see supplementary). To produce differently coloured prey items, we mixed the dough with black food colouring dye (Dr. Oetker, colour black, Bielefeld, Germany) for aposematically coloured prey, and green food dye (Dr. Oetker, colour green, Bielefeld, Germany) for the camouflaged version. For the aposematically coloured prey items, we painted three yellow dots using non-toxic yellow paint (PLUS Colour Acrylic Paint, Primary Yellow).

To create unpalatable prey items, we mixed bitrex solution (Stop & Grow, Mentholatum) into the dough at 7% concentration (see supplementary). Bitrex has been shown to make prey items unpalatable to great tits (Lindstedt et al., 2011), domestic chicks (Skelhorn & Rowe, 2005) and against invertebrate and vertebrate predators in the wild (Carroll & Sherratt, 2013; McLellan et al., 2022) in previous studies. Prey items were 2 cm long and 0.5 cm wide. To ensure that camouflaged prey with or without bitrex did not reflect significant amounts of ultraviolet wavelengths, potentially making them highly conspicuous to birds, we tested the composition of the dough for UV reflectance by using a spectrometer (Ocean Insight, model HR4PRO-XR-ES) with Oceanview software. In general, all prey items exhibited very low UV reflectance (**Fig. S2-S6**).

Altogether, we established eight treatment groups to summarize all potential combinations of defence strategies (**Table 1**): 1) solitary, camouflaged and palatable; 2) solitary, camouflaged and unpalatable; 3) solitary, aposematic and palatable; 4) solitary, aposematic and unpalatable; 5) grouped, camouflaged and palatable; 6) grouped, camouflaged and unpalatable; 7) grouped, aposematic and palatable; and 8) grouped, aposematic and unpalatable. Prey items were exposed to predators in the field from trays (8.5 cm diameter) constructed following protocols outlined by Nokelainen *et al*. (2014) (see **Fig.S1**). Trays exhibited very low UV reflectance similar to camouflaged prey (**Fig S7**).

Trays containing prey items were deployed along seven transects, each ∼2km in length, in three field locations in southern and central Finland (**Fig. S8**). Within each transect, we placed the trays 25 m apart from one another. The order of treatments within each transect was randomised beforehand using a lottery, and the same order was applied across all transects throughout the experiment. The predation experiment was conducted over two years. In 2023, we collected data from four transects located in Southern and Central Finland, with ten replicates per treatment per transect (i.e. 40 trays per treatment, 320 trays in total, resulting in 960 prey items). In 2024, we sampled from three transects located in Southern Finland with twelve replicates per treatment per transect (36 trays per treatment, 288 trays in total, resulting in a total of 864 prey items). Thus, in total, we had 76 trays for each of the eight treatment groups. Three trays were damaged during the experiment, leaving a total of 605 trays (with 1813 prey items overall).

Field predation experiments were conducted between late May and mid-June in both years (2023 and 2024) when Lepidopteran caterpillars are most abundant in Finland (Mappes et al., 2014). After the trays containing prey items were deployed in the field (day 1), we checked them once per day for attack marks or consumed prey (i.e., prey items that had disappeared from the tray) over the course of five days (days 2- 6). On the sixth day, after the attacks were recorded, all trays with prey items were collected from the field.

Prey was recorded as “attacked” if it had distinct bite marks or if it was partly or totally eaten (see **Fig. S9** for examples of attack marks). If the prey was not entirely consumed, we recorded whether the predator was an invertebrate or a bird based on the type of marks left on the tray. The attacked or missing prey items were replaced with new ones, or, if the mark was very minor (e.g. **Fig. S9D**), the surface was smoothed to maintain the odd ratios throughout the experiment (i.e., group treatments remained consistent as groups throughout the study).

### Invertebrate predator community

To measure the density and diversity of invertebrate predators along the study transects, we used pitfall traps buried in the ground (35 cl, 115 × 90 × 35 mm plastic containers filled with approximately 20 cl of water and fragrance-free detergent to reduce surface tension). The pitfall traps were arranged similarly to our forest heterogeneity measurements (see below). We placed the traps between every fifth and sixth tray, maintaining a distance of 10 m from each other throughout the experiment in both years. In some cases (e.g., if the ground was particularly rocky), the traps were located closer to the sixth tray.

In total, we placed 64 traps in 2023 (one transect had 19 traps, another had 13, while the remaining two transects each had 16 traps) and 60 in 2024 (20 traps per transect). Two traps were destroyed in 2024, resulting in a cumulative total of 122 traps. We set the traps on the second day of the experiment and emptied them on the fourth day. The insects from each trap were stored separately in plastic zip-lock bags in a -20°C freezer. From each sample, we recorded the number of individual invertebrates and identified them to the family level. Altogether, we collected 5210 individual invertebrate predators from 27 families (**Table S1**). We used the numbers of predatory invertebrates and number of families as covariates in our analyses.

We also recorded for each transect the abundance and number of bird species by using a standardized bird transect count method (in sensu Sutherland 2006) between 4 am and 8 am, on either the second or third day of the experiment depending on weather conditions (sunny and dry morning). The composition of the bird communities per transect are reported in **Table S1**. However, as the feeding territories of birds were likely to overlap between trays to some extent, and because we had only seven long transects in total due to the high number of treatments per transect, we were unable to reliably analyse the correlations between the insectivorous bird community per transect and predation risk, given that we only had seven data points (see, for example, Nokelainen et al., 2014 with 76 transects).

### Environmental variables

Due to logistic reasons, environmental variables were recorded from every tenth tray along each transect in 2023 (40 measurements) and every fifth tray in 2024 (58 measurements; see **Fig. S10** for a schematic of the sampling design). From these locations, we recorded the following measurements from a circular plot with a 5 m radius (with the tray located in the centre of the circle; see **Table S2** for details on how measurements were collected): 1) the number of living trees (i.e., tree density per plot); 2) the number of living tree species (see **Table S1** for the species list); 3) tree stem diameter at chest height (130 cm); 4) tree stem height (for trees with a diameter of at least 10 cm); 5) ground vegetation cover (percentage of trees and vegetation within the plot with diameters <10 cm); 6) basal area coverage of tree stands in m² per hectare (m^2^/ha); 7) proportion of dead trees per plot; 8) deadwood continuum; and 9) forest structure.

The latter two variables, deadwood continuum and forest structure, were used to define the natural state of the forest. The deadwood continuum (the decaying state of deadwood) was assessed using a refined visual assessment method for decaying wood (Tapio, 2023), where: 0 = no deadwood at all; 1 = weak deadwood continuum (the majority of the deadwood is fresh and composed of recently deceased standing trees); 2 = moderate deadwood continuum (the site contains a minimum of one-third of older and more decayed deadwood lying on the ground); and 3 = good deadwood continuum (the site contains at least one-third of deadwood from the previous categories among both standing and lying deadwood) (**Table S3**). For the statistical analyses, we combined deadwood continuum categories 0 and 1, as the number of category 0 instances was very low (66 out of 1813 plots). Similarly, there were only 43 plots belonging to category 3, which were combined with category 2 (485 plots). We refer to these two categories as weak and moderate-to-good deadwood continuum classes, with category 1 dominating the weak class (1219 out of 1285) and category 2 dominating the moderate-to-good class (485 out of 528).

Forest structure was estimated on a scale of 1 to 3 by following protocols from the 11th Finnish national forest inventory method (Valtakunnan Metsien Inventointi, VMI11, LUKE 2009). In this classification, category 1 indicates a naturally structured forest with randomly distributed trees of various age classes and species diversity (i.e., heterogenic forest structure), while category 3 denotes a human-managed, uniform, and homogeneous forest structure (see **Table S3** for more details). Category 2 falls between these two extremes, indicating a stand that has emerged naturally but has undergone an intermediate level of human management (e.g., some thinning and the removal of larger trees).

### Statistical analyses

To match environmental data to the trays sampled, we assumed that environmental variables collected in 2023 were similar for four trays before and five trays after the measurement point, and in 2024 for two trays before and after the measurement point (see **Fig. S10**).

To analyse whether the survival of prey varied between different antipredator defence treatments (hypotheses **I**-**IV**), we employed a linear mixed effects model with a binomial distribution and a logit function. The response variable was the initial attack risk (i.e., the first attack(s) per prey item), classified as either attacked or not attacked (model 1, **Table S4**). The fixed effects included prey colouration (camouflaged or aposematic), palatability (palatable or unpalatable), group size (solitary or grouped), and their three-way interactions. As random effects, we included year of the experiment and the transect ID nested within each year.

To analyse how the survival of prey items (irrespective of the antipredator defence treatment) varied based on environmental conditions (**V**), we conducted a linear mixed model analysis with a binomial distribution and logit function, using the initial attack risk of prey as a binomial response variable. As we were interested in how the predator community, vegetational structure and forest heterogeneity could impact survival of our prey items, we used three separate models (model 2-4, **Table S4**) for analysis.

In the first model (model 2, **Table S4**), we analysed how the number and diversity of predatory invertebrates influence the survival of prey. The number of invertebrate predators and family taxon groups were included as continuous explanatory variables in the model.

In the second model (model 3, **Table S4**), we analysed how vegetation structure influences survival. The number of tree species, individual number of trees, tree height, tree diameter, ground vegetation cover, basal area cover and proportion of deadwood were included as continuous explanatory variables in this model. Before conducting the analysis, we examined how the variables covaried through two-way correlations using the cor.test function in R version 4.4.1 (R Core Team, 2024) and by visually examining plotted data. Significant correlations were found among all variables; however, the coefficients of most were weak and did not appear to correlate when visually inspected (**Fig S8**). Therefore, we decided to consider only correlations with coefficients greater than 0.5 as meaningful, as these appeared to show actual correlations when plotted. Under these conditions we found correlations between tree height and tree diameter, as well as between the number of individual trees and ground vegetation cover (see **Fig S8**). Consequently, the final versions of the model excluded tree stem diameter and the number of individual trees.

In the third model (model 4, **Table S4**), we examined how survival was affected by categorical variables describing the level of natural state and the deadwood continuum in the forest. The forest structure categories (three categories) and deadwood continuum (two categories) were included as fixed variables. Additionally, we analysed whether the number of invertebrate predators and the number of invertebrate predator families differed between deadwood continuum categories and forest structure categories (models 5-6, **Table S4**).

Finally, to test for interactive effects between the antipredator defence treatment and environmental conditions (**VI**), we employed the same set of environmental variables as in models 2-4 in three new separate models (models 8-10, **Table S4**). We also included prey colour, palatability, and gregariousness as fixed variables, along with two-way interactions between the antipredator defence components and environmental variables. For all our analyses we used the glmmTMB package version 1.1.10 (Brooks et al., 2017) in R version 4.4.1 (R Core Team, 2024) with logit function, except for model 6, which utilised the lme4 package 1.1-35.5 (Bates et al., 2015) due to the data being normally distributed. For pairwise comparisons of forest structure with three categories, we employed the emmeans package version 1.10.5 (Lenth, 2024). Model assumptions were confirmed with the DHARMA package version 0.4.7 (Hartig, 2024) using the simulateResiduals function to test model assumptions, testDispersion for overfitting and testZeroInflation for zero-inflation.

## Results

### Gregariousness increases the survival of the prey

Of the 605 trays, 179 (30%) were attacked at least once (**Table 1**). The attack rates by all predators (invertebrates and birds) varied between 22% and 35% across all experimental days (**Table 1**). Survival of solitary prey was significantly lower compared to that of gregarious prey (**Table 2**, **Fig. 1**). Colouration (camouflaged vs. aposematic) or palatability (palatable vs. unpalatable) did not significantly affect survival (**Table 2**, **Fig. 1**). There were no significant two-way or three-way interactions between the antipredator defence treatments regarding survival (**Table 2**).

**Fig 1.**
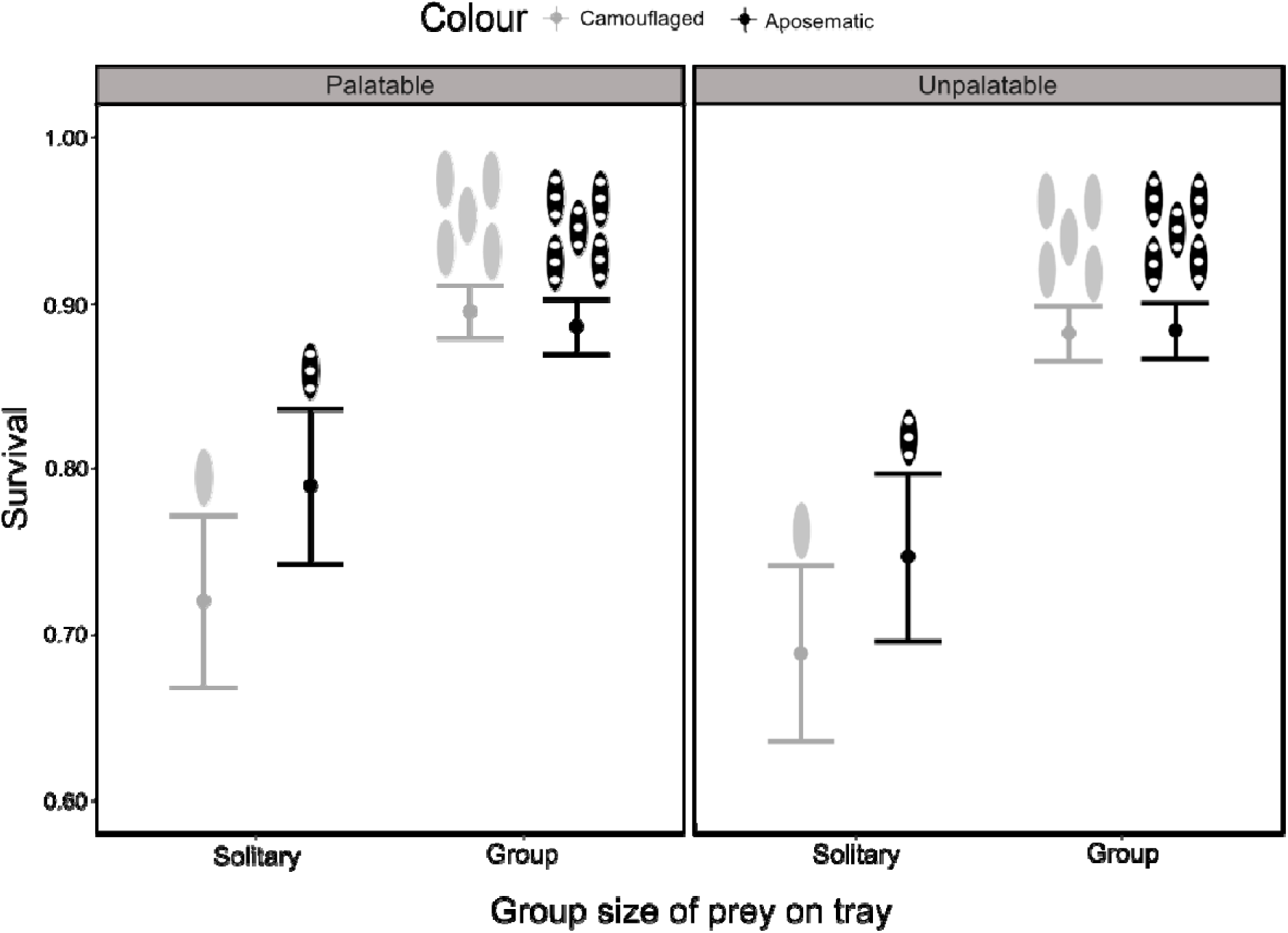
Effect of antipredator defence on survival of prey items: group size (solitary or in groups of five), colouration (“camouflaged” green & “aposematic” black and yellow) and palatability (Palatable & unpalatable with bitrex).

**Table 2.**
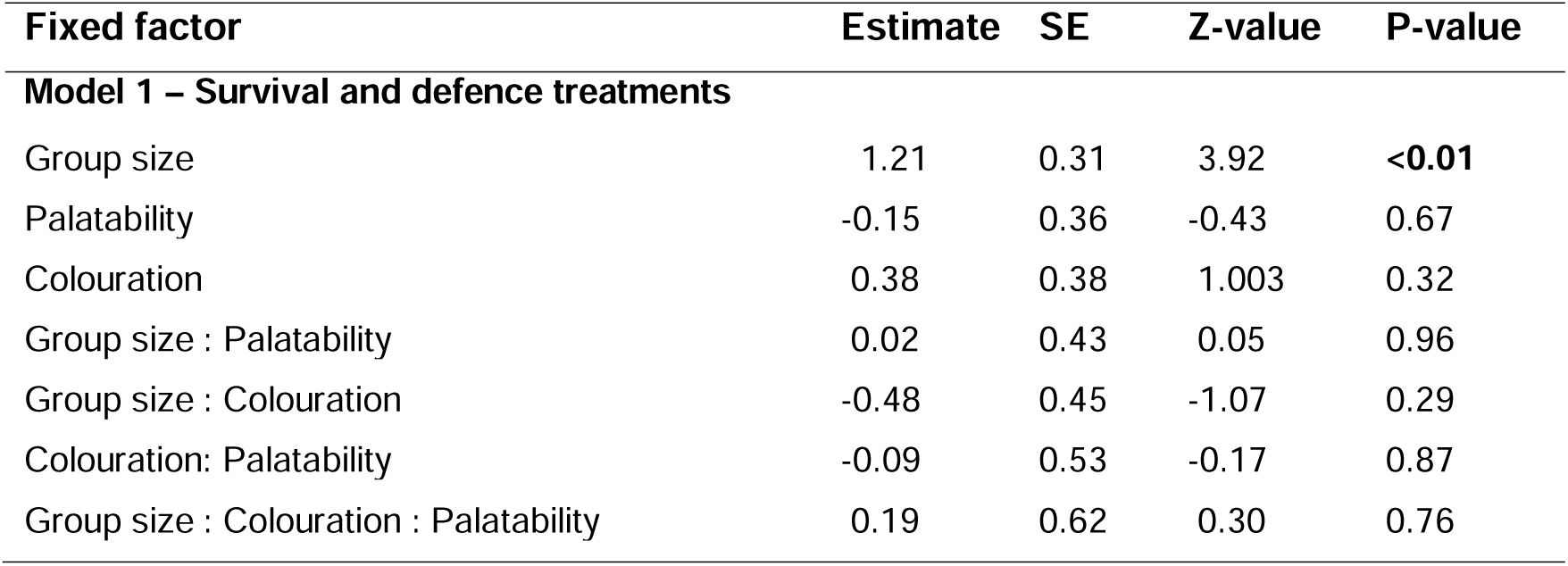
Summary table of model 1 analysing survival based on antipredator defence treatments. Effects for the dependent variables were compared against “undefended” equivalents (I.e., group size = “solitary”; palatability = “palatable”; colouration = “camouflaged”). Significant p-values (p<=0.05) are marked in bold.

| **Fixed factor** | **Estimate** | **SE** | **Z-value** | **P-value** |
|---|---|---|---|---|
| **Model 1 – Survival and defence treatments** |  |  |  |  |
| Group size | 1.21 | 0.31 | 3.92 | **<0.01** |
| Palatability | -0.15 | 0.36 | -0.43 | 0.67 |
| Colouration | 0.38 | 0.38 | 1.003 | 0.32 |
| Group size : Palatability | 0.02 | 0.43 | 0.05 | 0.96 |
| Group size : Colouration | -0.48 | 0.45 | -1.07 | 0.29 |
| Colouration: Palatability | -0.09 | 0.53 | -0.17 | 0.87 |
| Group size : Colouration : Palatability | 0.19 | 0.62 | 0.30 | 0.76 |

### Survival was lower in a more heterogenous environment

The local abundance of invertebrate predators and the number of invertebrate predator families did not have a significant effect on the survival of prey (**Table 3**). Tree species number, mean tree height, ground vegetation cover and the percentage of deadwood per study plot did not significantly affect survival (**Table 3**). However, prey items were more likely to be attacked in areas with a higher basal area cover (i.e. a higher density of trees per hectare) (**Table 3**).

**Table 3.**
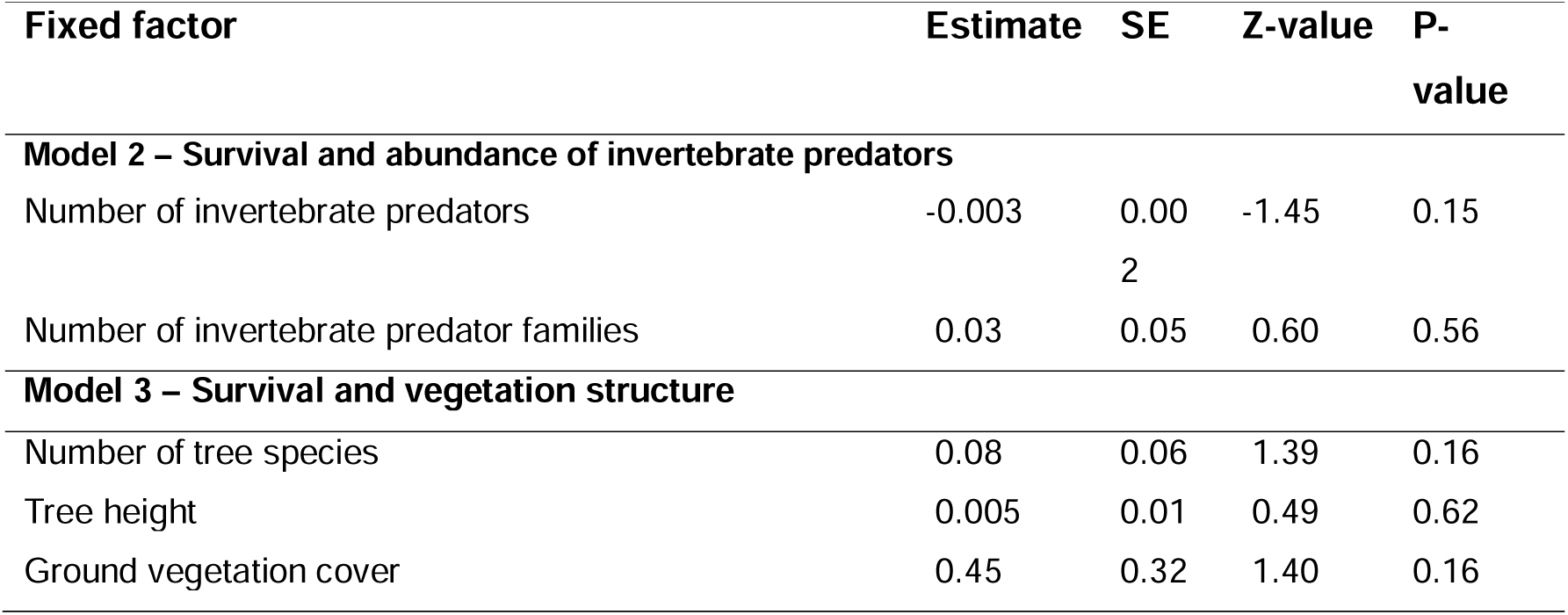
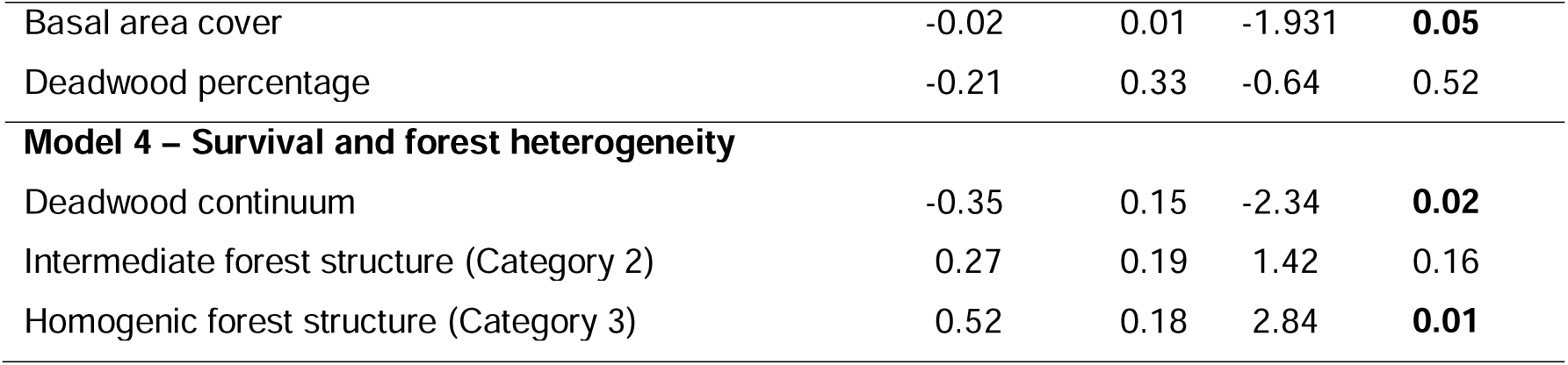
Summary table of models 2-4 analysing survival based on environmental variables. Significant p-values (p<=0.05) are marked in bold.

| Fixed factor | Estimate | SE | Z-value | P-value |
|---|---|---|---|---|
| **Model 2 – Survival and abundance of invertebrate predators** |  |  |  |  |
| Number of invertebrate predators | -0.003 | 0.002 | -1.45 | 0.15 |
| Number of invertebrate predator families | 0.03 | 0.05 | 0.60 | 0.56 |
| **Model 3 – Survival and vegetation structure** |  |  |  |  |
| Number of tree species | 0.08 | 0.06 | 1.39 | 0.16 |
| Tree height | 0.005 | 0.01 | 0.49 | 0.62 |
| Ground vegetation cover | 0.45 | 0.32 | 1.40 | 0.16 |
| Basal area cover | -0.02 | 0.01 | -1.931 | **0.05** |
| Deadwood percentage | -0.21 | 0.33 | -0.64 | 0.52 |
| **Model 4 – Survival and forest heterogeneity** |  |  |  |  |
| Deadwood continuum | -0.35 | 0.15 | -2.34 | **0.02** |
| Intermediate forest structure (Category 2) | 0.27 | 0.19 | 1.42 | 0.16 |
| Homogenic forest structure (Category 3) | 0.52 | 0.18 | 2.84 | **0.01** |

When we analysed the impact of deadwood continuum and forest structure on prey survival, we found that survival was significantly lower (i.e., predation pressure was higher) in areas with a moderate-to-good deadwood continuum compared to areas where the deadwood continuum was categorised as weak (**Table 3**, **Fig. 2A**). Similarly, survival was significantly lower in areas where the forest structure resembled a natural state (category 1) compared to areas with a homogeneous, human-managed structure (category 3) (Estimate = -0.52, p = 0.01). There were no significant differences in survival between forest structure categories 1 (natural state or similar to natural) and 2 (intermediate heterogeneity) (Estimate = -0.27, p = 0.33), or between category 2 and category 3 (homogeneous) (Estimate = -0.25, p = 0.38).

**Fig 2.**
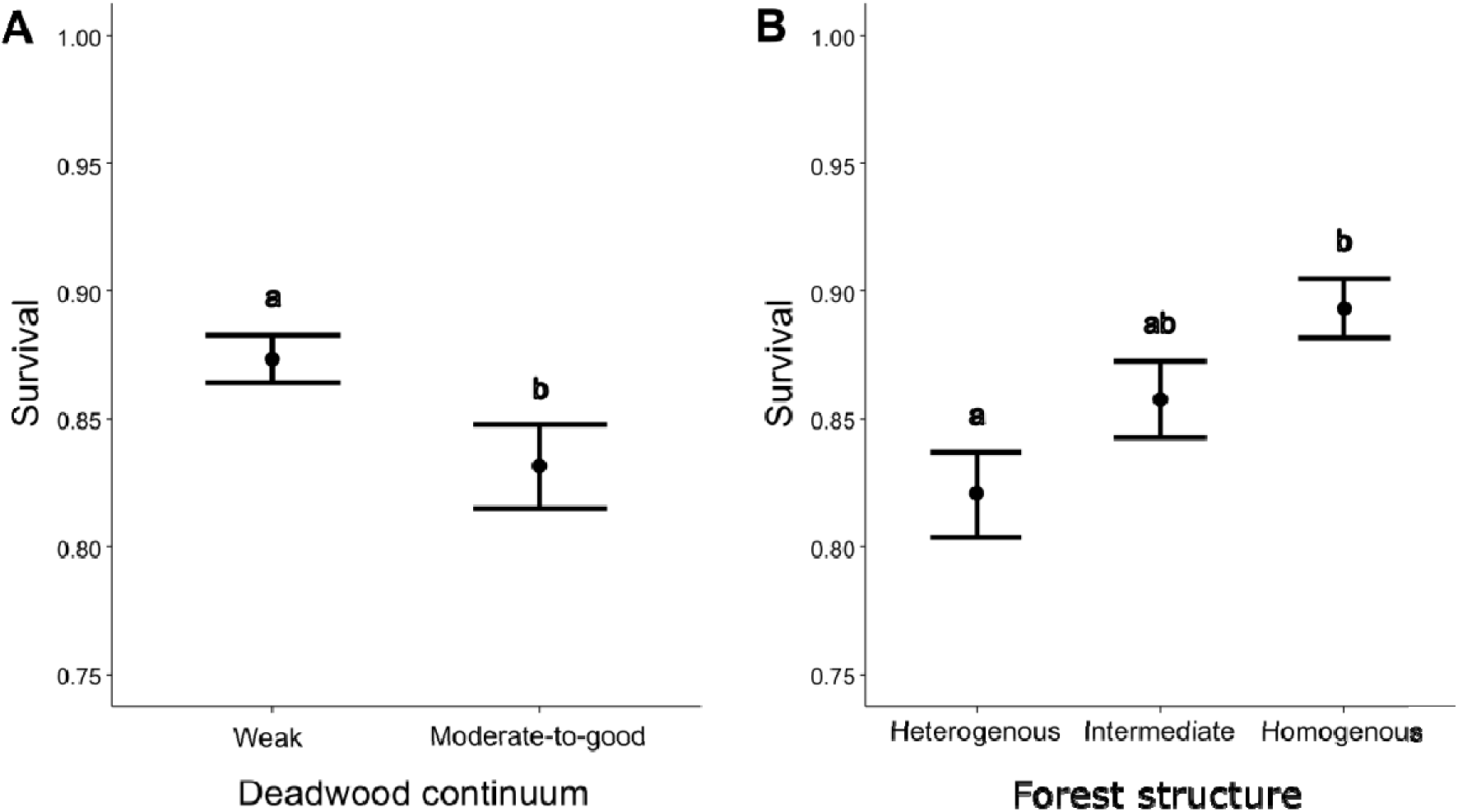
Effect of forest heterogeneity categories on survival of artificial prey: **A)** Survival in two categories of deadwood continuum (weak deadwood continuum = deadwood is either missing or the majority is fresh and composed from the recently died standing trees; moderate-to-good deadwood continuum = minimum of 1/3 of older and more decayed deadwoods either standing or laying down). **B)** Survival in three categories of forest structure. Heterogenous (category 1) forest structure indicates a natural state forest with randomly distributed trees of various age classes and species diversity, with intermediate (category 2) and homogenous (category 3) meaning increasingly more human managed plots, respectively. See **Table S3** for more details.

Upon further examination of how the abundance of invertebrate predators varied under different forest structure categories, we found that the number of invertebrate predators was significantly higher in the more heterogenous natural state areas (category 1) compared to the homogeneously structured forest areas (category 3) (Estimate = 0.31, p = 0.02, **Fig 3A**), but did not differ between forest structure categories 1 (natural state or similar to natural) and 2 (intermediate heterogeneity) (Estimate = 0.11, p = 0.74), or between category 2 and category 3 (homogeneous) (Estimate = 0.20, p = 0.35). The number of invertebrate predator family taxa did not differ significantly based on the forest structure (category 1 vs. 2: Estimate = 0.27, p = 0.62; category 1 vs. 3: Estimate = 0.01, p = 0.99; category 2 vs. 3: Estimate = - 0.26, p = 0.58, **Fig 3B**). Finally, invertebrate predator abundance did not differ significantly between deadwood continuum categories (Estimate = 0.02, p = 0.89, **Fig 3C**) and neither did invertebrate predator families (Estimate = -0.24, p = 0.22, **Fig 3D**).

**Fig 3.**
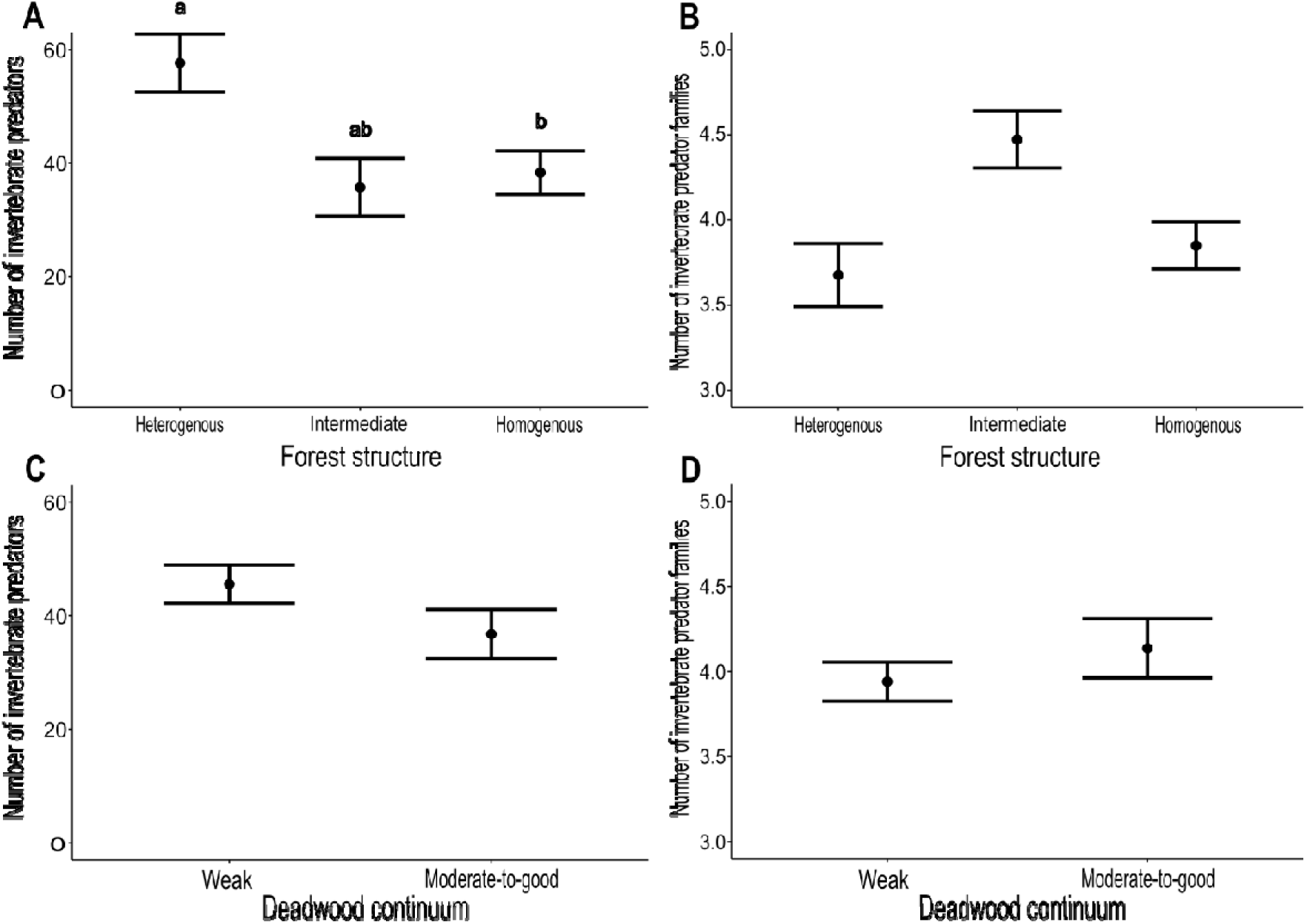
Invertebrate predator abundance and diversity across forest heterogeneity categories. **A)** Number of invertebrate predators across forest structure categories. **B)** Number of invertebrate predator families across forest structures. **C)**. Number of invertebrate predators across deadwood continuum categories. **D)** Number of invertebrate predator families across deadwood continuums.

### Survival rates of camouflaged and aposematic prey varied depending on environmental conditions

There were no significant interactions between the survival of any of the defence treatments (colour, palatability, or gregariousness) and the number of invertebrate predators (**Table 4**). Furthermore, there were no significant differences in the survival of prey among any of the defence treatments related to the number of invertebrate predator taxa (i.e. families, **Table 4**). Once again, gregariousness significantly increased the survival of prey irrespective of the invertebrate predator density or diversity (**Table 4**).

**Table 4.**
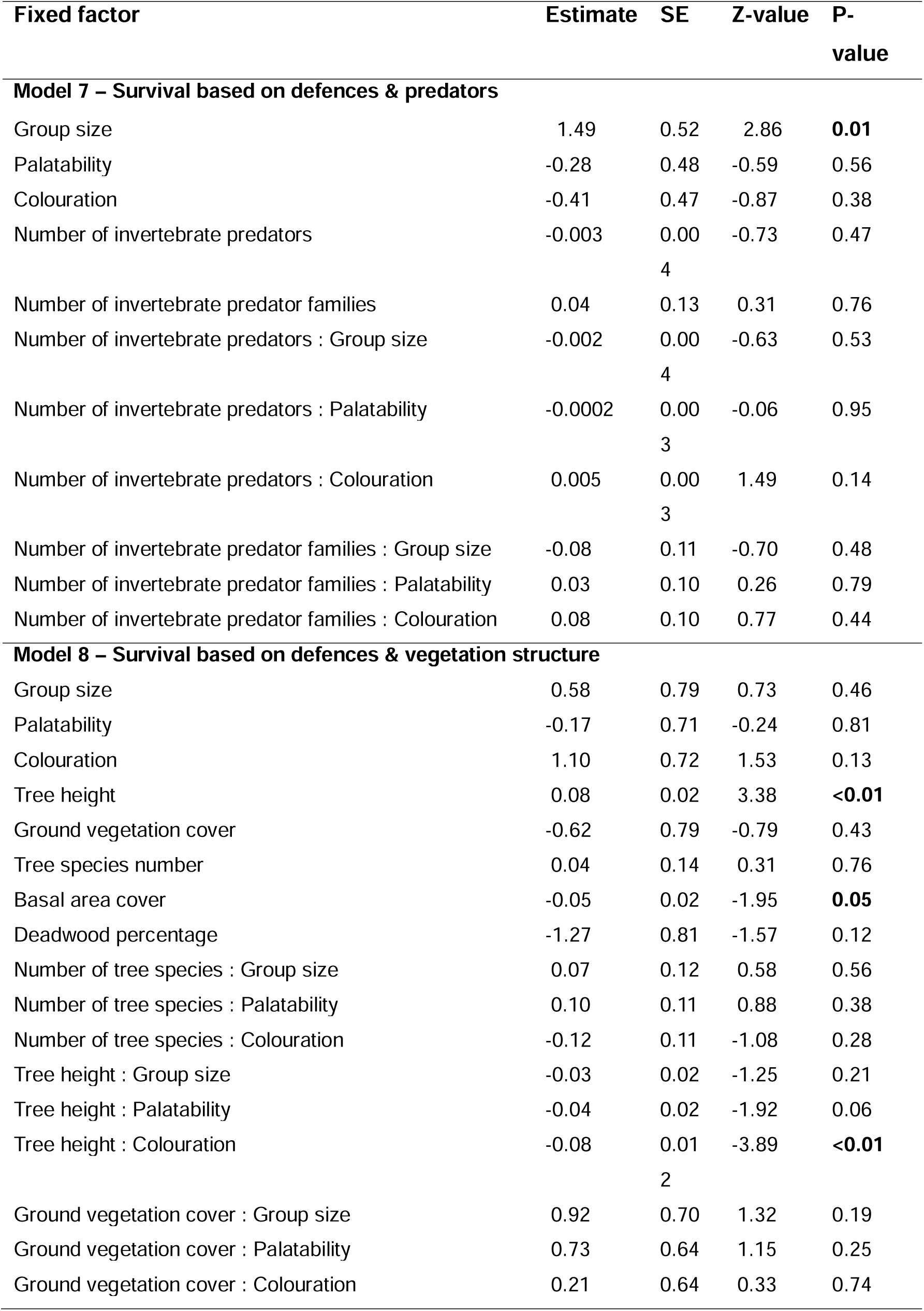
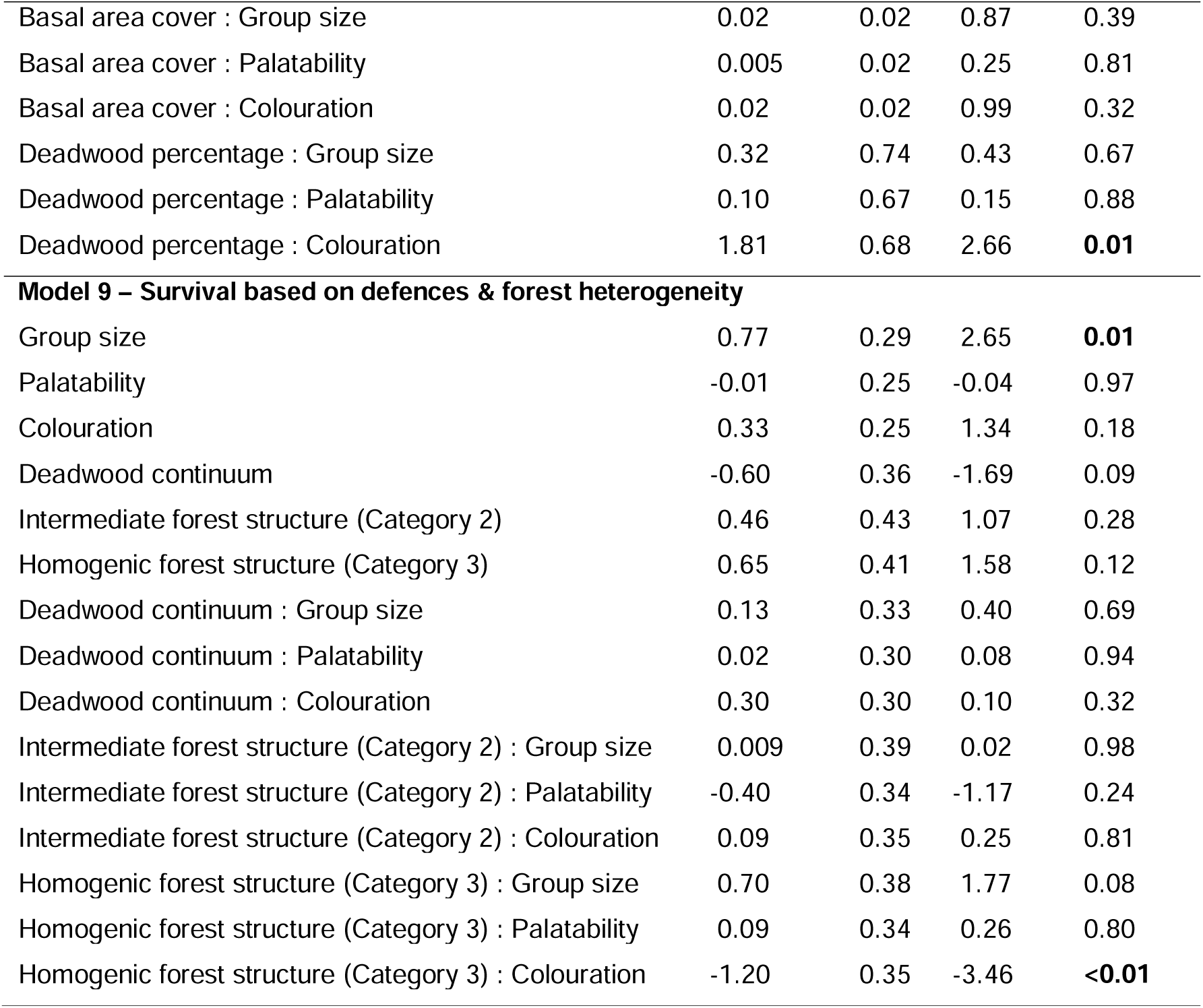
Summary table of model 7-9 analysing survival based on two-way interactions between antipredator defence treatments and environmental conditions.

| Fixed factor | Estimate | SE | Z-value | P-value |
|---|---|---|---|---|
| **Model 7 – Survival based on defences & predators** |  |  |  |  |
| Group size | 1.49 | 0.52 | 2.86 | **0.01** |
| Palatability | -0.28 | 0.48 | -0.59 | 0.56 |
| Colouration | -0.41 | 0.47 | -0.87 | 0.38 |
| Number of invertebrate predators | -0.003 | 0.004 | -0.73 | 0.47 |
| Number of invertebrate predator families | 0.04 | 0.13 | 0.31 | 0.76 |
| Number of invertebrate predators : Group size | -0.002 | 0.004 | -0.63 | 0.53 |
| Number of invertebrate predators : Palatability | -0.0002 | 0.003 | -0.06 | 0.95 |
| Number of invertebrate predators : Colouration | 0.005 | 0.003 | 1.49 | 0.14 |
| Number of invertebrate predator families : Group size | -0.08 | 0.11 | -0.70 | 0.48 |
| Number of invertebrate predator families : Palatability | 0.03 | 0.10 | 0.26 | 0.79 |
| Number of invertebrate predator families : Colouration | 0.08 | 0.10 | 0.77 | 0.44 |
| **Model 8 – Survival based on defences & vegetation structure** |  |  |  |  |
| Group size | 0.58 | 0.79 | 0.73 | 0.46 |
| Palatability | -0.17 | 0.71 | -0.24 | 0.81 |
| Colouration | 1.10 | 0.72 | 1.53 | 0.13 |
| Tree height | 0.08 | 0.02 | 3.38 | **<0.01** |
| Ground vegetation cover | -0.62 | 0.79 | -0.79 | 0.43 |
| Tree species number | 0.04 | 0.14 | 0.31 | 0.76 |
| Basal area cover | -0.05 | 0.02 | -1.95 | **0.05** |
| Deadwood percentage | -1.27 | 0.81 | -1.57 | 0.12 |
| Number of tree species : Group size | 0.07 | 0.12 | 0.58 | 0.56 |
| Number of tree species : Palatability | 0.10 | 0.11 | 0.88 | 0.38 |
| Number of tree species : Colouration | -0.12 | 0.11 | -1.08 | 0.28 |
| Tree height : Group size | -0.03 | 0.02 | -1.25 | 0.21 |
| Tree height : Palatability | -0.04 | 0.02 | -1.92 | 0.06 |
| Tree height : Colouration | -0.08 | 0.012 | -3.89 | **<0.01** |
| Ground vegetation cover : Group size | 0.92 | 0.70 | 1.32 | 0.19 |
| Ground vegetation cover : Palatability | 0.73 | 0.64 | 1.15 | 0.25 |
| Ground vegetation cover : Colouration | 0.21 | 0.64 | 0.33 | 0.74 |
| Basal area cover : Group size | 0.02 | 0.02 | 0.87 | 0.39 |
| Basal area cover : Palatability | 0.005 | 0.02 | 0.25 | 0.81 |
| Basal area cover : Colouration | 0.02 | 0.02 | 0.99 | 0.32 |
| Deadwood percentage : Group size | 0.32 | 0.74 | 0.43 | 0.67 |
| Deadwood percentage : Palatability | 0.10 | 0.67 | 0.15 | 0.88 |
| Deadwood percentage : Colouration | 1.81 | 0.68 | 2.66 | **0.01** |
| **Model 9 – Survival based on defences & forest heterogeneity** |  |  |  |  |
| Group size | 0.77 | 0.29 | 2.65 | **0.01** |
| Palatability | -0.01 | 0.25 | -0.04 | 0.97 |
| Colouration | 0.33 | 0.25 | 1.34 | 0.18 |
| Deadwood continuum | -0.60 | 0.36 | -1.69 | 0.09 |
| Intermediate forest structure (Category 2) | 0.46 | 0.43 | 1.07 | 0.28 |
| Homogenic forest structure (Category 3) | 0.65 | 0.41 | 1.58 | 0.12 |
| Deadwood continuum : Group size | 0.13 | 0.33 | 0.40 | 0.69 |
| Deadwood continuum : Palatability | 0.02 | 0.30 | 0.08 | 0.94 |
| Deadwood continuum : Colouration | 0.30 | 0.30 | 0.10 | 0.32 |
| Intermediate forest structure (Category 2) : Group size | 0.009 | 0.39 | 0.02 | 0.98 |
| Intermediate forest structure (Category 2) : Palatability | -0.40 | 0.34 | -1.17 | 0.24 |
| Intermediate forest structure (Category 2) : Colouration | 0.09 | 0.35 | 0.25 | 0.81 |
| Homogenic forest structure (Category 3) : Group size | 0.70 | 0.38 | 1.77 | 0.08 |
| Homogenic forest structure (Category 3) : Palatability | 0.09 | 0.34 | 0.26 | 0.80 |
| Homogenic forest structure (Category 3) : Colouration | -1.20 | 0.35 | -3.46 | **<0.01** |

In terms of vegetation, there were significant interactions between prey colouration and tree height, as well as between prey colouration and the percentage of dead wood (**model 8, Table 4**). Specifically, the survival of camouflaged prey was higher in taller tree stands (**Fig. 4A**) and when the percentage of dead wood was low (**Fig. 4B**), in comparison to aposematically coloured prey. Besides basal area cover, other vegetation variables had no significant main effects on the survival of different prey types (**Table 4**).

**Fig 4.**
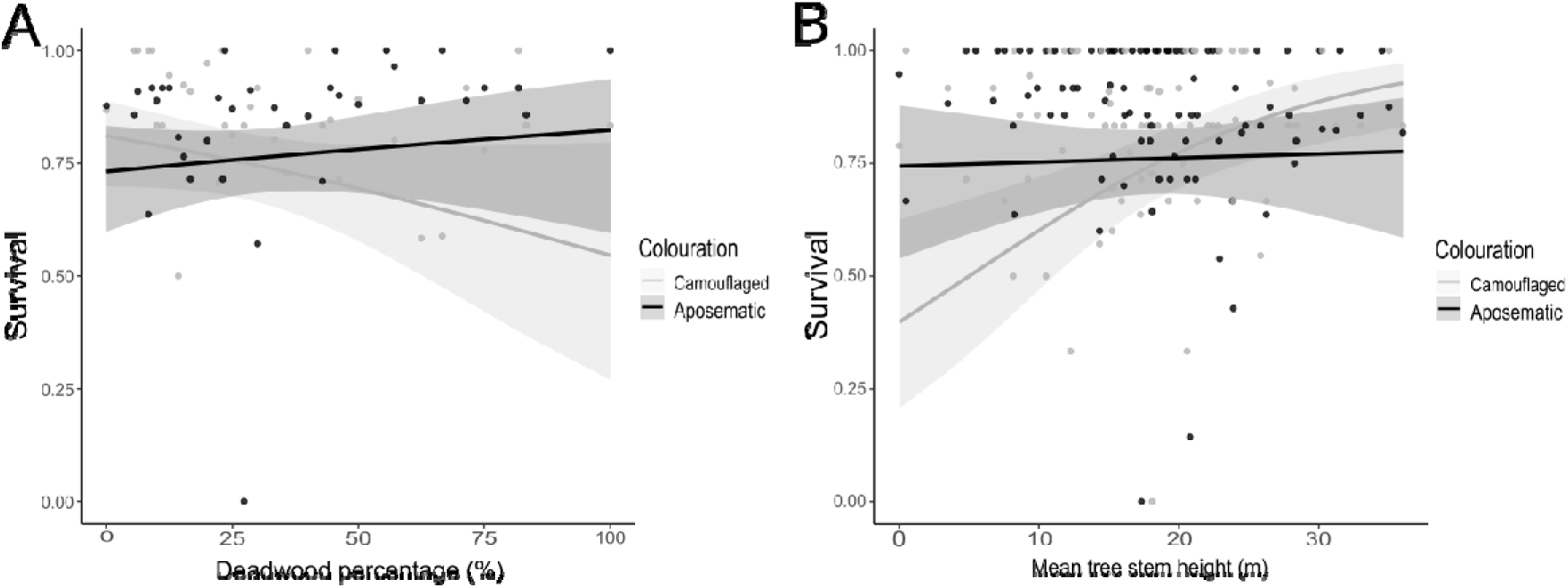
Interactive effects of antipredator defence treatments and environmental conditions on the survival of artificial prey. **A)** Two-way interaction between deadwood percentage and colouration. **B)** Two-way interaction between tree stem height and colouration.

There was a significant interaction between prey colouration and forest structure category 3 (**Table 4**). Specifically, aposematic prey had similar survival across all forest structure categories (category 1 vs. 2: Estimate = -0.35, p = 0.46; category 1 vs. 3: Estimate = 0.17, p = 0.79; category 2 vs. 3: Estimate = 0.53, p = 0.15), while camouflaged prey were attacked more in heterogenous environments compared to areas with more managed structure (category 1 vs. 3: Estimate = -1.03, p < 0.01; category 2 vs. 3: Estimate = -0.76, p = 0.02). There was no significant difference in survival of camouflaged prey between heterogenous and intermediately heterogenous forest structure areas (category 1 vs 2: Estimate = -0.27, p = 0.57) (**Fig. 5**). There were no significant interactions between the deadwood continuum categories and antipredator defence strategies (**Table 4**). Once again, prey individuals in groups exhibited significantly higher overall survival rates, irrespective of environmental conditions (**Table 4**).

**Fig 5.**
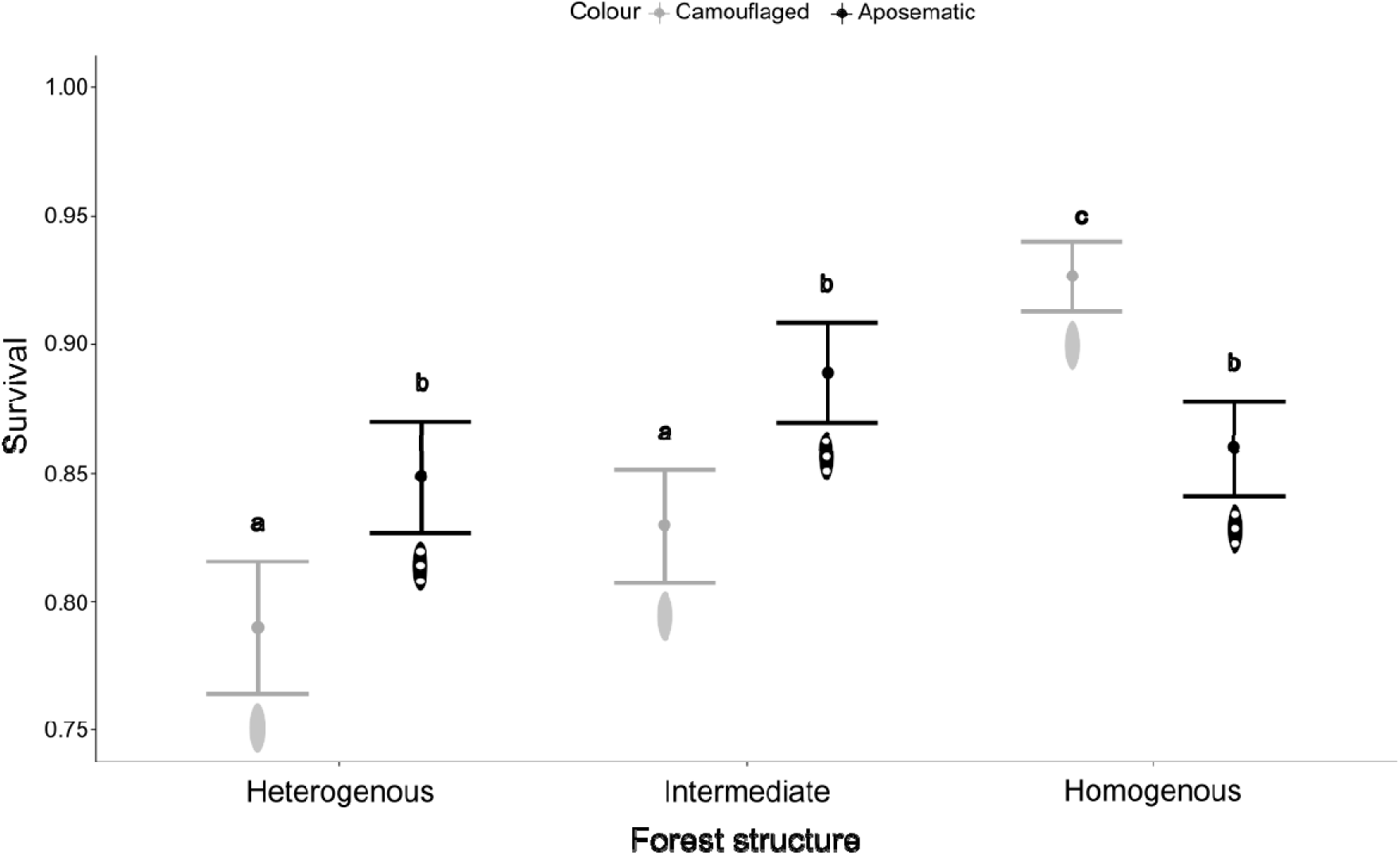
Interactive effects of colouration treatment and forest structure categories on the survival of artificial prey.

## Discussion

To better understand the factors affecting the invasion success of pest insects in novel environments, we need to determine mechanisms that improve their survival against predators. Here we demonstrate that gregariousness could be one of the key traits as it increased survival of our artificial prey items against naturally varying predator communities. Interestingly, the impact of aposematic colouration, unpalatability, and their various combinations, had no independent effects on survival. Our results also suggest that canopy cover, diversity in forest structure, and a high-quality deadwood continuum increased predation pressure, thereby potentially making heterogenic forests more resilient to pest invasions. This could partly be attributed to the higher numbers of invertebrate predators found in more heterogeneously structured forests.

Gregariousness during the larval stage is often associated with the outbreak tendency in herbivorous insects (Battisti et al., 2023; Hunter, 2000). In line with this, our results suggest that decreased predation risk can be one of the mechanisms underlying this phenomenon (Ronnås et al., 2010; Reynolds et al., 2009). The survival was higher in groups, and we did not detect any indications of an increased detectability risk for grouped prey (but see: Lindstedt et al., 2006, 2011). We also found no evidence to suggest that other defensive strategies (i.e., unpalatability and warning colours) had a strong effect on the survival of individuals in groups. This contrasts with some other empirical studies, which have found gregariousness to be beneficial for aposematically coloured prey but not for cryptic prey (Beatty et al. 2005; McLellan et al. 2022). Instead, our results indicate that group living can reduce per capita mortality risk, mainly through dilution effects and/or predator satiation (McClure & Despland, 2011; Riipi et al., 2001; Van Meyel et al., 2026), making gregariousness beneficial also for the cryptically coloured prey in terms of survival against predators. Consequently, when applied to forest pest insects, gregariousness may provide and enemy-free time-window for population growth in a new environment, facilitating invasion into new areas.

Efficient chemical defences and aposematism have sometimes been associated with the success of invasive species, such as the lantern fly (Song et al. 2018), fire ants (Wickings & Ruberson 2016) and cane toads (Shine 2010). However, in our study, we found no differences in survival between aposematically coloured and camouflaged prey, nor between unpalatable and palatable prey. One explanation for this finding is that the predators in the field likely varied extensively in their sensitivity for the manipulated defensive traits (see e.g. Mappes & Endler, 2004; Rowland et al., 2017, Kikuchi et al., 2021). For example, while defensive colouration plays an important role in protection against avian predators (Hossie & Sherratt, 2013; Mappes et al. 2014; Rönkä et al. 2020), ants and predatory beetles may be less discriminative when attacking warningly coloured prey (Dyer 1995; Sugiura 2020; Yilmaz & Spaethe 2022). Considering this natural variation among predators and its effects on predation risk is therefore very important as when potential pest insects invade a new area, they will encounter not only visually foraging avian predators but also various other predator species, such as ants, spiders, and predatory beetles, all likely to vary in their responses to the visual and chemical defence characteristics of the prey (Kikuchi et al., 2021).

It is crucial to acknowledge that our study design and surveys of predator community structure present certain limitations. First, while our measurements of predation pressure can be generalised to reflect the attack risk imposed by generalist predators, they do not elucidate the roles of specialized predators and parasitoids specific to certain insect species, nor the mortality they may inflict (Duan et al., 2015). However, the predation pressure imposed by generalist predators, such as insectivorous birds, ants or predatory beetles, has been suggested to play an important role in determining the potential for insect populations to begin growing exponentially at the onset of an outbreak (Elkinton et al., 1996; Mason et al., 1983; Parry et al., 1997), while specialists such as parasitoids and diseases are more influential in terminating outbreaks (Dwyer et al. 2004; Snyder & Ives 2003). Therefore, when predicting the factors that influence invasion success in various environments, the community of generalist predators and their propensity to attack particular prey are in a key role (Kenis et al., 2017).

Second, our survey of the predator community is not comprehensive. While the use of edible artificial prey allowed us to manipulate and test the effects of unpalatability on predation risk, we were unable to reliably identify the full range of predators attacking the models through bite marks. This is because many prey items were consumed entirely by one or more predators, leaving no discernible beak or bite marks to analyse (Barnett et al., 2016; Hossie & Sherratt, 2013). This contrasts with experiments that employed clay models, which are often tasted but not wholly consumed, allowing for a greater number of bite marks to assess the predator community (Galappaththige et al., 2024; Kollberg et al., 2022; Lawrence et al., 2019). Additionally, our measurement of the invertebrate predator community relied on pitfall traps, which primarily capture non-flying invertebrate predators and fail to collect flying invertebrate predators, such as dragonflies, wasps and robber flies. Thus, in future studies, it would be important to quantify the insectivorous predator community more broadly as well as measure the amount of alternative prey available for predators and how it correlates with the predation pressure on the focal preys.

Third, while bitrex has been shown to render prey items unpalatable to birds (Skelhorn & Rowe, 2005; 2010) and impart a remarkably bitter taste to humans (M. Jonsson, pers. obs.), it may not have been equally distasteful to all potential predators attacking the prey in the field. The concentration of bitrex in the dough was a bit higher (7%) than that used in some other studies (often between 0.5 – 2.5%) (Lindstedt et al., 2011; McLellan et al., 2022; Carroll & Sherratt, 2013). Thus, in the light of our results the manipulation was likely comparable to a mildly unprofitable chemical defence when estimated against the whole natural predator community. However, if invasive species deploy highly toxic or deterrent chemical defences, e.g. similar to those of cane toads (Hayes et al., 2009) or fire ants (Long et al., 2015), it most likely enhances survival and facilitates dispersal to new areas.

Previous studies have demonstrated that forests with higher tree species diversity and age structures often correlate with higher abundance and diversity of predators and parasitoids (Bellone et al., 2017; Bereczki et al., 2014; Blomqvist et al., 2016; Kollberg et al., 2022; Rappa et al., 2024: Wrethling et al., 2026). Our results align with these studies, suggesting that both predation pressure and the abundance of invertebrate predators are greater in heterogeneous forests. Specifically, we observed that predation pressure was higher in environments characterized by dense canopy covers, a moderate-to-good deadwood continuum, and a more complex and natural forest structure. While we did not find a direct association between the abundance of invertebrate predators and predation risk, we did find that the number of invertebrate predators was higher in more natural forests compared to managed and homogeneous ones (**Fig 3**), alongside elevated attack rates in those areas (**Fig. 2**, **Table 3**). Moreover, our findings indicate that, rather than the sheer amount of deadwood, the quality of the deadwood continuum is associated with a higher predation pressure. Other studies have reported similar results, highlighting that the quality rather than the volume of deadwood increases local invertebrate predator diversity (Seibold et al., 2016; Similä et al., 2003).

We did not find support for our hypothesis that predation risk for aposematic prey would increase with forest heterogeneity due to a higher density and diversity of potential predators. Conversely, the predation risk for aposematic prey varied less across different environmental variables. Significant interactions were observed between canopy height, the amount of deadwood, and forest structure, primarily driven by changes in the survival of camouflaged prey. The greater variation in survival linked to environmental conditions in camouflaged prey may indicate that variations in the visual environment are more critical for the efficacy of camouflage compared to warning colours, similar to findings from a recent global scale study (Medina et al., 2025). Furthermore, variation in predator community structure or predator densities may influence attack rates for camouflaged and aposematic prey along these environmental gradients (Medina et al., 2025; Nokelainen et al., 2014). Finally, our analyses of the interactive effects of environment and defence strategy consistently showed that grouped prey had higher survival rates, suggesting that gregariousness has a stronger impact on survival, irrespective of environmental conditions or other defensive strategies employed.

In conclusion, our study emphasizes the importance of considering species ecology such as gregariousness and how it is combined with different defensive strategies when predicting the success of invasive pest insects. This should also be taken into account in population dynamic models applied to estimate outbreak risks as predator-induced mortality is rarely static, but rather varies depending on the prey’s defensive strategy. Finally, our findings contribute to the growing body of literature suggesting that protecting and maintaining heterogeneity in economically managed forests could enhance their resilience against insect outbreaks (Blomqvist et al. 2016; Bellone et al., 2017; Bereczki et al., 2014; Kollberg et al., 2014; 2022; Root, 1973; Russell 1989; Wrethling et al., 2026). This approach can reduce outbreak risks through multiple mechanisms, including the preservation of higher densities of generalist predators that regulate the population densities of forest pest insects (Bereczki et al., 2014; Eveleigh et al., 2007; Nixon & Roland, 2012), as well as decreasing the likelihood of pest insects finding suitable host plants (Kollberg et al., 2022; Hambäck & Englund, 2005). Increasing the habitat heterogeneity and different aged deadwood in economically managed forests would also benefit many endangered forest species (Huuskonen et al., 2021; Hekkala et al., 2023; Heidrich et al., 2023; Cuff et al., 2021).

## Ethical note

BitrexTM is non-toxic and does not harm any potential predator species when ingested. Experiments were conducted in accordance with the ethical guidelines of University of Helsinki (TENK).

## Author contribution statement

V.K., M.J., K.K. and C.L. designed research; V.K., M.J. and K.K. performed research; M.J., V.K. and E.G. analysed data; and all authors wrote the paper.

## Acknowledgements

This study was funded by the Research Council of Finland via the project no 330578 (C.L.) and 333482 (C.L.), by the University of Helsinki doctoral programme of wildlife biology (LUOVA) (M.J.), by the Finnish Cultural Foundation via a research grant no 00220400 (M.J.) and 00240558 (M.J.) and supported by the Humboldt fellowship programme for postdocs (E.G). We would like to thank Martti Sarkia and Kimmo Peltola for their help in the lab. We would also like to thank Tarmo Ketola for their assistance.

