## Supplemental Figures and Tables for "Gregariousness can promote invasion success of pest insects while higher environmental heterogeneity decreases it"

**Supplementary materials**

**S1. Study system**

Artificial prey items used in the experiment where made by mixing wheat flour (Coops Xtra) with vegetable butter (Sunnuntai munkinpaistorasva, a mixture of palm, coconut, and canola oil; manufacturer: Bunge Finland Oy) in a 2:1 ratio. For the camouflaged colour treatment, we mixed the dough with food colouring dye (50 ml per 100 g of dough), using green food dye (Dr. Oetker, colour black, Bielefeld, Germany). For the aposematic colour treatment, we mixed the dough black food dye (Dr. Oetker, colour green, Bielefeld, Germany) using the same ratio as for camouflaged. In addition to the black colour, we painted three yellow dots using non-toxic yellow paint (PLUS Colour Acrylic Paint, Primary Yellow) on the aposematic prey items after they had been shaped into caterpillars. All prey items were stored in cold temperatures (8°C) while not being used in the field.

Trays where prey items were located consisted of a petri dish (8.5cm in diameter) painted green (two layers of Tikkurila, M384 “Basilika green”) glued onto flower sticks using water-soluble, odourless superglue (Loctite Super Glue Creative). We used rust-free, non-shimmering nuts, combined with paper tape and metal washers, to stabilise the sticks to the petri dishes. The height of the trays was 90 cm (**Fig. S1**).

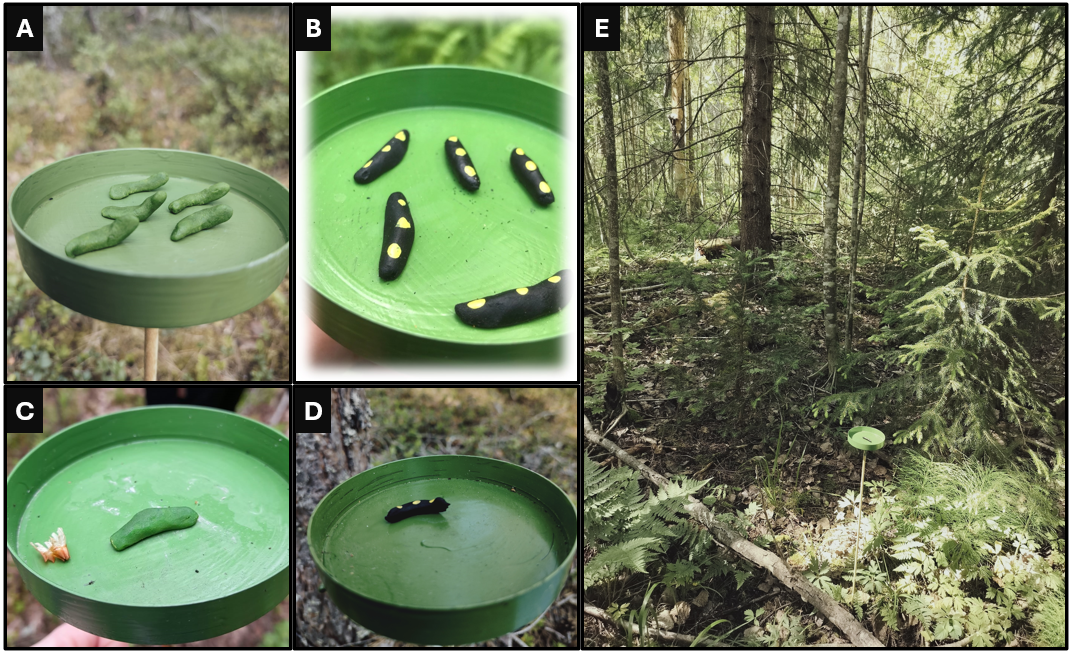

**Fig S1**. Artificial prey items used for measuring predation risk. **A)** Grouped and camouflaged prey. **B)** Grouped and aposematic. **C)** Solitary and camouflaged. **D)** Solitary and aposematic. **E)** Tray placed in the forest. Height of trays was 90cm above ground. Dimensions of a prey item were 2x0.5 cm. Photos A, B, D, and E were taken by Magnus Jonsson. Photo C was taken by Milja Lipponen (reproduced with permission).

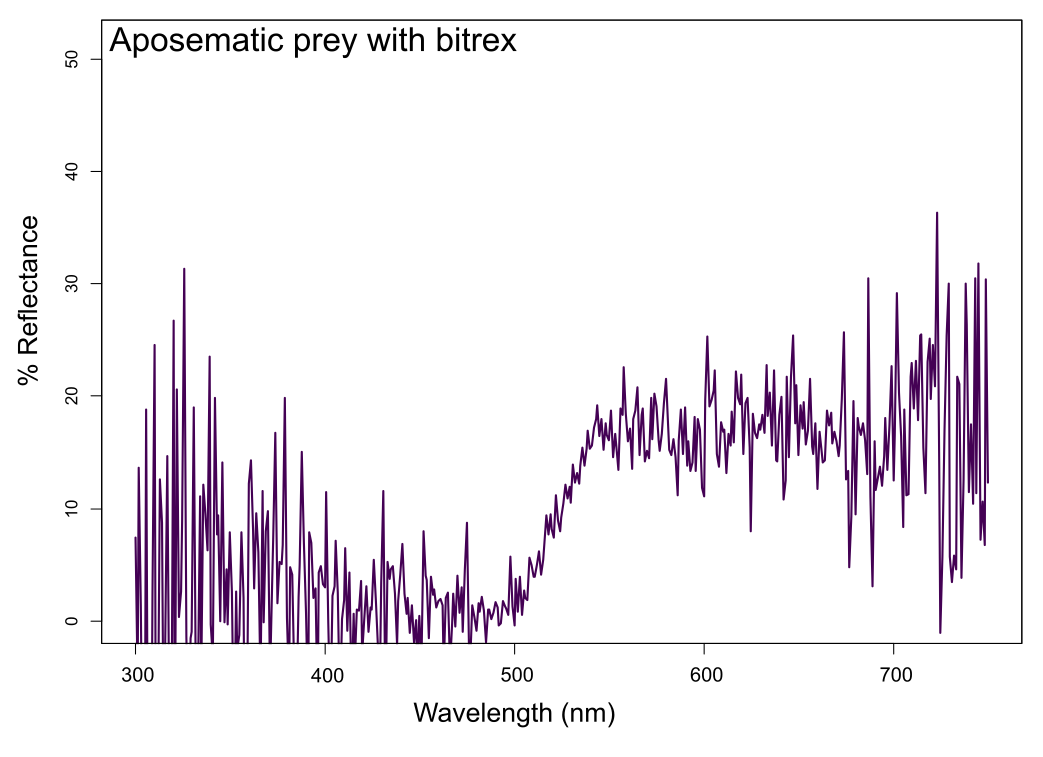

**Fig S2.** Spectral reflectance (%) of artificial prey dough containing black food dye (Dr. Oetker) and bittering agent (Bitrex; Stop & Grow, Mentholatum), measured between 300 and 750 nm. A high reflectance in the wavelengths for UV light (300-400) is an indication that our prey items would be conspicuous towards birds or other predators who can spot these wavelengths.

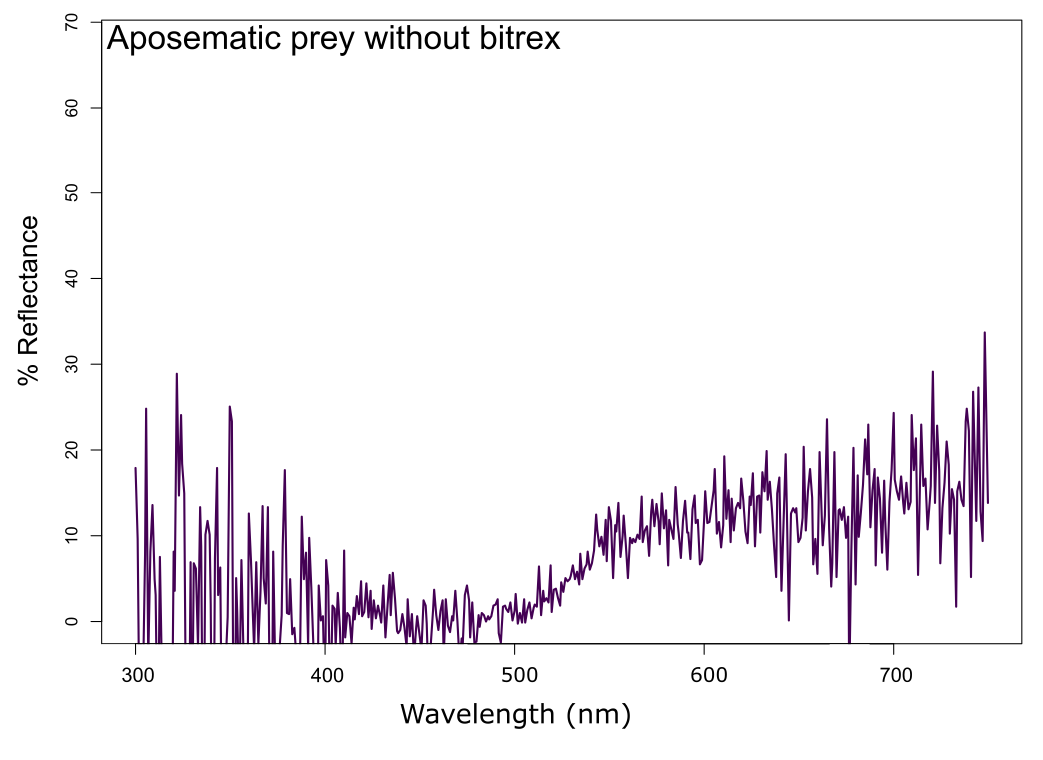

**Fig S3.** Spectral reflectance (%) of artificial prey dough containing black food dye, measured between 300 and 750 nm.

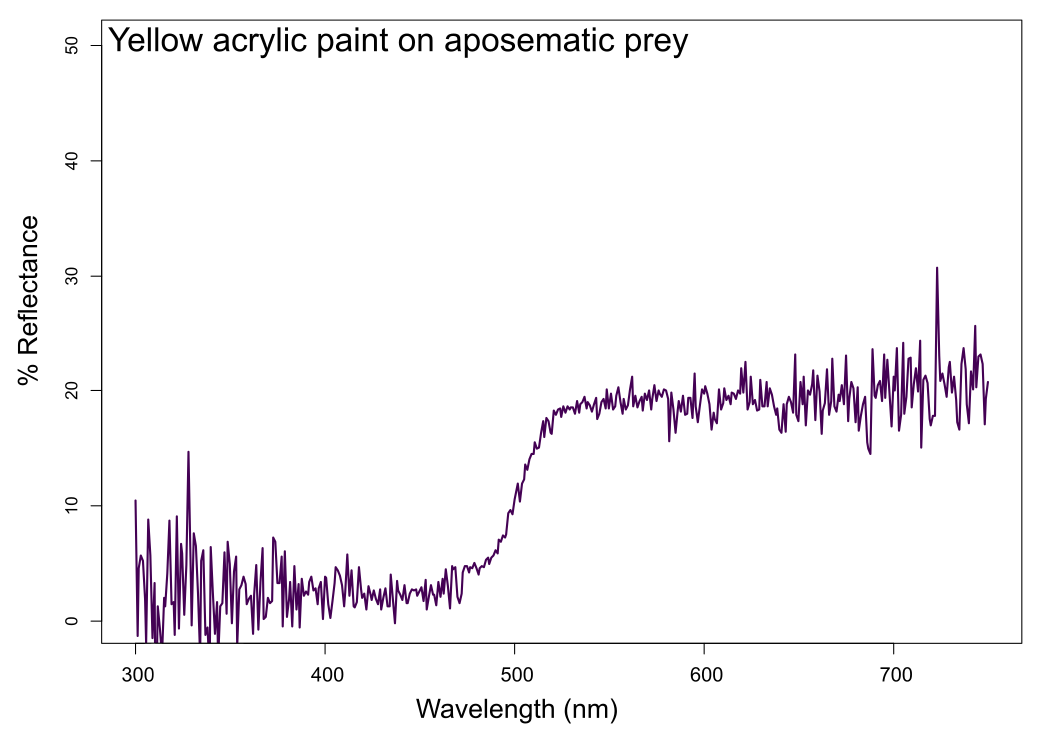

**Fig S4**. Spectral reflectance (%) of yellow acrylic paint (PLUS colour acrylic paint Primary yellow) used on aposematic (black) prey, measured between 300 and 750 nm.

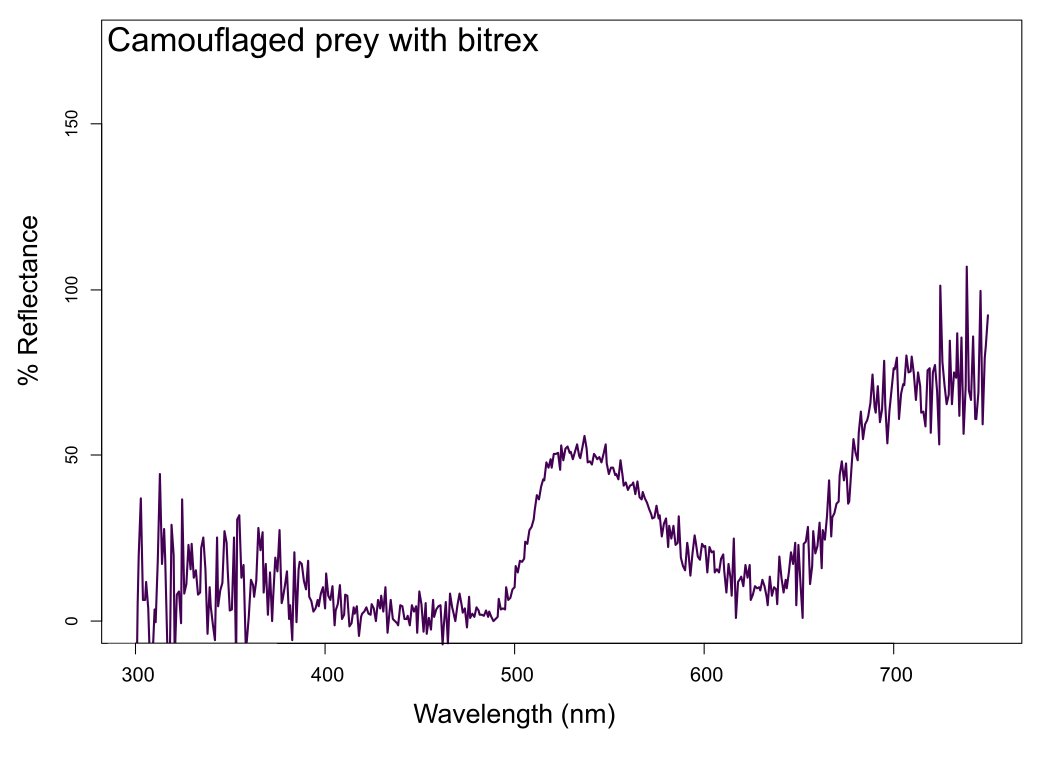

**Fig S5**. Spectral reflectance (%) of artificial prey dough containing green food dye and bittering agent, measured between 300 and 750 nm.

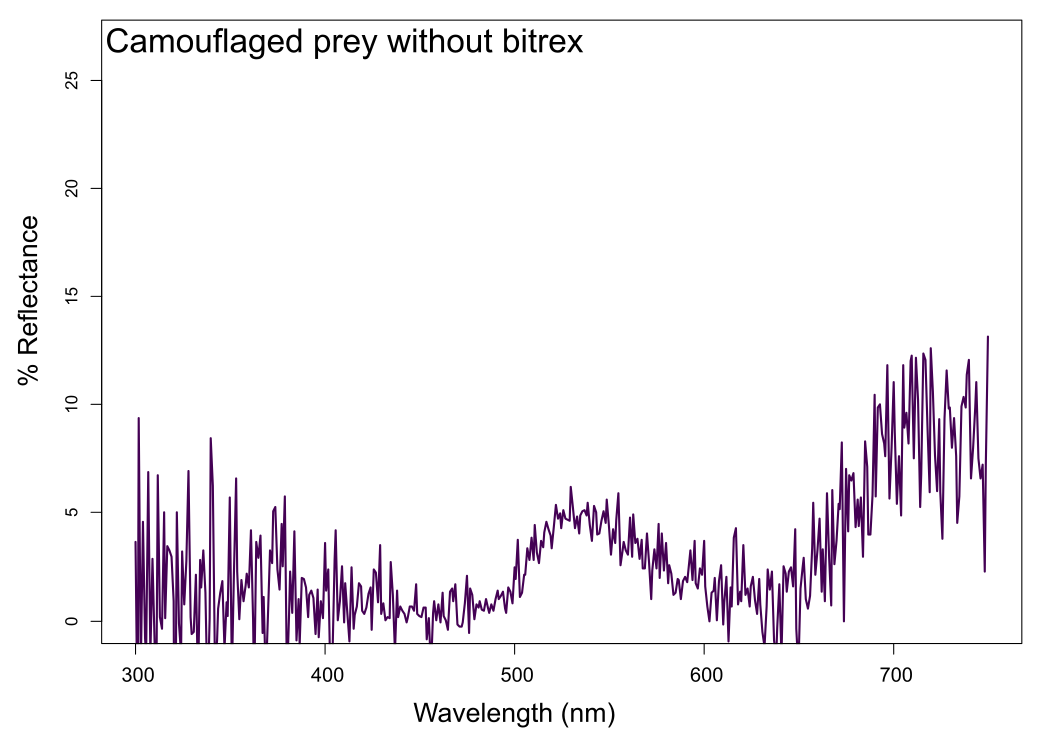

**Fig S6.** Spectral reflectance (%) of artificial prey dough containing green food dye, measured between 300 and 750 nm.

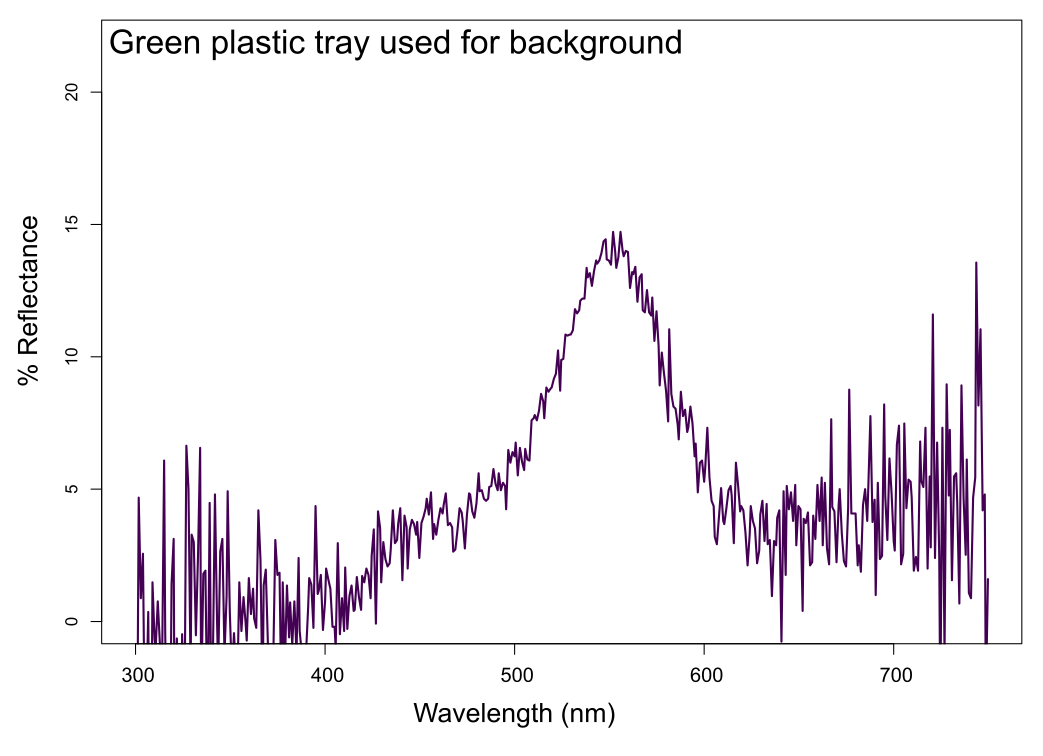

**Fig S7**. Spectral reflectance (%) of tray used as background for prey during the experiment, painted with green paint (Tikkurila, M384 “Basilika green”), measured between 300 and 750nm.

**S2. Data collection**

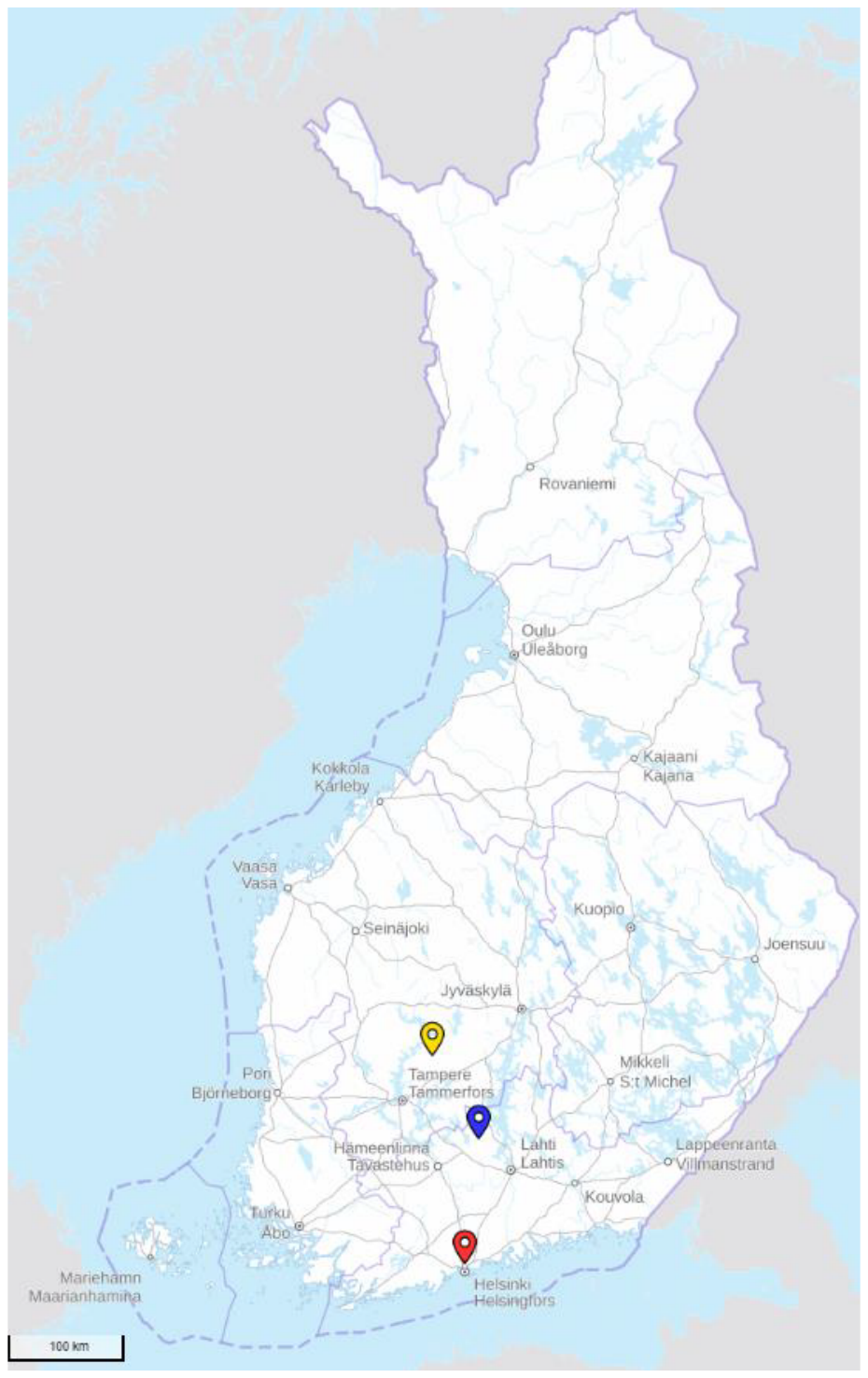

**Fig S8**. Map of Finland and locales where transects took place during 2023 and 2024. Altogether, we had two transects in Hyytiälä (the most northern point; 61.84519402550942, 24.28731921207365; 61.83524453993747, 24.311799383372946), one in Evo (middle point; 61.19295583168848, 25.108410563685368) and four in Helsinki (the most southern point; two in Maunula forest [60.22439497621672, 24.91931393008953, 60.2211957927351, 24.916871514426344]; one in Paloheinä [60.26139561787009, 24.913931723156285] and one in Viikki arboretum [60.21725826109837, 25.03048359922209]), resulting in seven transects. Transects in Hyytiälä were done in 2023. Transect in Evo was done in 2024. Two of the transects in Helsinki (Maunula 1 and 2) were done in 2023 and the other two (Paloheinä and Viikki) in 2024. Three trays were destroyed during the experiment from the transect in Paloheinä (Helsinki) and were excluded from all analyses. Two pitfall traps were lost in Evo and were also excluded from the analysis on invertebrate numbers (marked as NA in model 3, 6 and 7, see **Table s4**). Background map © Maanmittauslaitos (2024).

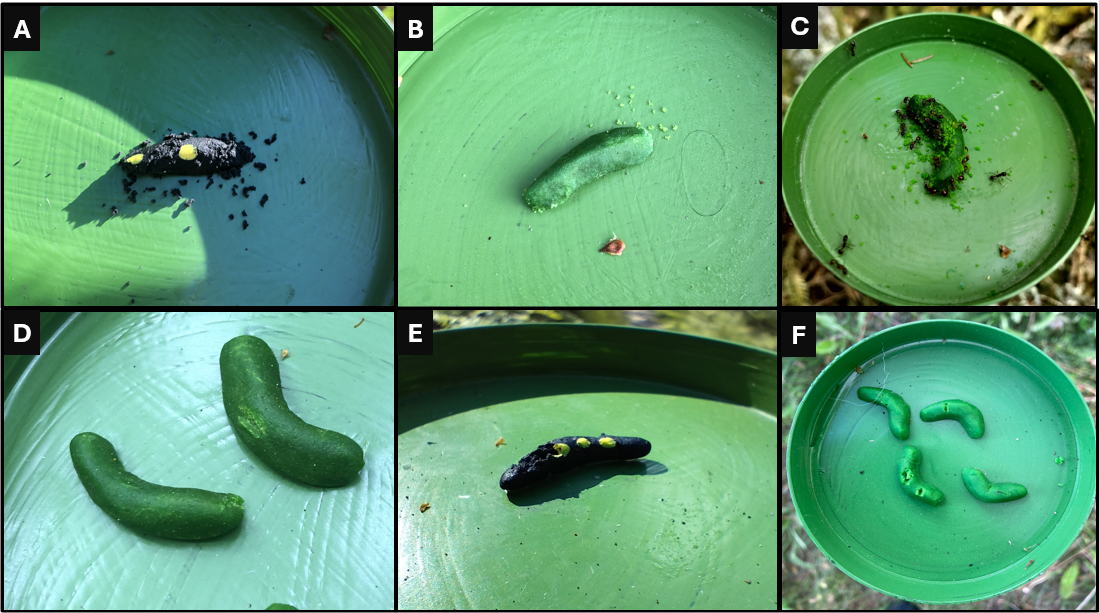

**Fig S9**. Examples of attack marks on artificial prey items. **A-B)** Examples of invertebrate attack marks. **C)** Ants attacking artificial prey. **D-F)** Examples of bird attack marks. Definitions of attacks were made using definitions of previous, similar studies (e.g. Schwab et al., 2020; Low et al., 2014). Photos A, B and D were taken by Venla Korhonen. Photos C, E and F were taken by Magnus Jonsson.

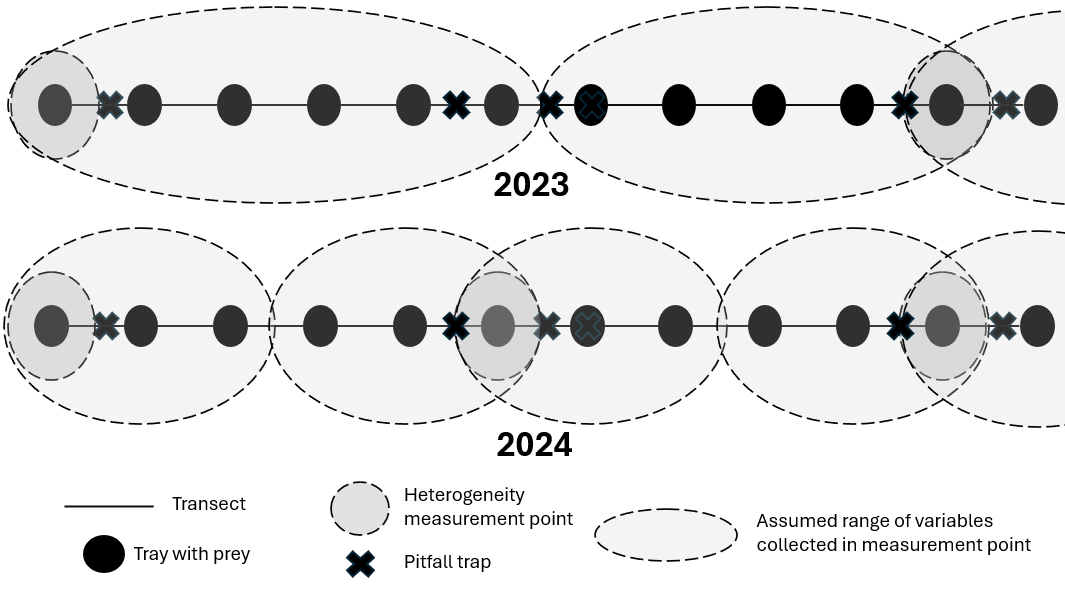

**Fig S10.** Illustration showcasing sampling design for environmental variables. In 2023, due to logistic limitations, we sampled environmental data from every 10^th^ tray and assumed that each measured point represented the next 5 trays and subsequent 4 trays along the transect. In 2023, we sampled every 5^th^ site and assumed each measured point represented the following 2 trays and subsequent 2 trays along the transect. We placed pitfall traps to collect data on the invertebrate predator community after every 5^th^ tray in both years.

**Table S1.** Transect specific species (or family) lists of birds, trees and invertebrates collected in 2023 and 2024.

| **Transect** | **Bird species ID** | **Tree species ID** | **Invertebrate family ID** |
| --- | --- | --- | --- |
| **2023** |  |  |  |
| **Maunula 1 (Helsinki)** | *Certhia familiaris* | *Alnus incana* | *Carabidae* |
|  | *Chloris chloris* | *Betula sp.* | *Formicidae* |
|  | *Corvus corone cornix* | *Betula pendula* | *Lithobiidae* |
|  | *Cyanistes caeruleus* | *Betula pubescens* | *Lycosidae* |
|  | *Dendrocopos major* | *Juniperus sp.* | *Phalangidae* |
|  | *Dryobates minor* | *Picea abies* | *Pisauridae* |
|  | *Emberiza citrinella* | *Pinus sylvestris* | *Thomisidae* |
|  | *Erithacus rubecula* | *Populus tremula* | *Tetragnathidae* |
|  | *Ficedula hypoleuca* | *Salix aurita* | *Staphylinidae* |
|  | *Fringilla coelebs* | *Sorbus aucuparia* |  |
|  | *Motacilla alba* | *Quercus sp.* |  |
|  | *Muscicapa striata* |  |  |
|  | *Parus major* |  |  |
|  | *Periparus ater* |  |  |
|  | *Phylloscopus sibilatrix* |  |  |
|  | *Phylloscopus trochilus* |  |  |
|  | *Pyrrhula pyrrhula* |  |  |
|  | *Regulus regulus* |  |  |
|  | *Spinus spinus* |  |  |
|  | *Sylvia atricapilla* |  |  |
|  | *Troglodytes troglodytes* |  |  |
|  | *Turdus iliacus* |  |  |
|  | *Turdus merula* |  |  |
|  | *Turdus philomelos* |  |  |
| Sum | 133 | 84 | 1385 |
| **Maunula 2 (Helsinki)** | *Certhia familiaris* | *Acer sp.* | *Araneidae* |
|  | *Chloris chloris* | *Alnus incana* | *Carabidae* |
|  | *Corvus corax* | *Betula pendula* | *Formicidae* |
|  | *Chroicocephalus ridibundus* | *Betula pubescens* | *Lycosidae* |
|  | *Curruca communis* | *Picea abies* | *Phalangiidae* |
|  | *Cyanistes caeruleus* | *Pinus sylvestris* | *Philodromidae* |
|  | *Dendrocopos major* | *Populus tremula* | *Pisauridae* |
|  | *Erithacus rubecula* | *Prunus padus* | *Trombidiidae* |
|  | *Ficedula hypoleuca* | *Sorbus aucuparia* | *Tetragnathidae* |
|  | *Fringilla coelebs* | *Quercus sp.* | *Staphylinidae* |
|  | *Larus canus* |  |  |
|  | *Motacilla alba* |  |  |
|  | *Muscicapa striata* |  |  |
|  | *Parus major* |  |  |
|  | *Phylloscopus sibilatrix* |  |  |
|  | *Phylloscopus trochilus* |  |  |
|  | *Regulus regulus* |  |  |
|  | *Spinus spinus* |  |  |
|  | *Sylvia atricapilla* |  |  |
|  | *Sylvia borin* |  |  |
|  | *Troglodytes troglodytes* |  |  |
|  | *Turdus merula* |  |  |
|  | *Turdus philomelos* |  |  |
|  | *Turdus pilaris* |  |  |
| Sum | 145 | 67 | 742 |
| **Hyytiälä 1** | *Anthus trivialis* | *Betula sp.* | *Araneidae* |
|  | *Certhia familiaris* | *Betula pendula* | *Asilidae* |
|  | *Dendrocopos major* | *Betula pubescens* | *Carabidae* |
|  | *Erithacus rubecula* | *Picea abies* | *Clubionidae* |
|  | *Fringilla coelebs* | *Pinus sylvestris* | *Formicidae* |
|  | *Parus major* | *Populus tremula* | *Lithobiidae* |
|  | *Phoenicurus phoenicurus* | *Sorbus aucuparia* | *Lycosidae* |
|  | *Phylloscopus collybita* |  | *Phalangiidae* |
|  | *Phylloscopus trochilus* |  | *Philodromidae* |
|  | *Prunella modularis* |  | *Pisauridae* |
|  | *Regulus regulus* |  | *Salticidae* |
|  | *Spinus spinus* |  | *Sicariidae* |
|  | *Troglodytes troglodytes* |  | *Zoropsidae* |
|  | *Turdus merula* |  | *Trombidiidae* |
|  |  |  | *Thomisidae* |
|  |  |  | *Tetragnathidae* |
|  |  |  | *Staphylinidae* |
|  |  |  | *Spheciformes* |
| Sum | 63 | 67 | 625 |
| **Hyytiälä 2** | *Anthus trivialis* | *Alnus incana* | *Carabidae* |
|  | *Certhia familiaris* | *Betula sp.* | *Formicidae* |
|  | *Chloris chloris* | *Betula pendula* | *Lycosidae* |
|  | *Dendrocopos major* | *Betula pubescens* | *Phalangiidae* |
|  | *Erithacus rubecula* | *Picea abies* | *Philodromidae* |
|  | *Fringilla coelebs* | *Pinus sylvestris* | *Pisauridae* |
|  | *Garrulus glandarius* | *Populus tremula* | *Salticidae* |
|  | *Larus canus* | *Salix sp.* | *Trombidiidae* |
|  | *Lophophanes cristatus* | *Salix raita* | *Tetragnathidae* |
|  | *Parus major* | *Sorbus aucuparia* | *Staphylinidae* |
|  | *Phylloscopus collybita* |  |  |
|  | *Phylloscopus sibilatrix* |  |  |
|  | *Phylloscopus trochilus* |  |  |
|  | *Spinus spinus* |  |  |
|  | *Sylvia atricapilla* |  |  |
|  | *Sylvia borin* |  |  |
|  | *Troglodytes troglodytes* |  |  |
| Sum | 57 | 66 | 792 |
| **2024** |  |  |  |
| **Evo** | *Anthus trivialis* | *Alnus incana* | *Blattoidea* |
|  | *Corvus cornix* | *Betula pubescens* | *Carabidae* |
|  | *Columba palumbus* | *Juniperus communis* | *Elateridae* |
|  | *Dendrocopos major* | *Picea abies* | *Formicidae* |
|  | *Erithacus rubecula* | *Pinus sylvestris* | *Gnaphosidae* |
|  | *Ficedula hypoleuca* | *Populus tremula* | *Histeridae* |
|  | *Fringilla coelebs* | *Salix raita* | *Linyphiidae* |
|  | *Parus major* | *Sorbus aucuparia* | *Lycosidae* |
|  | *Phoenicurus phoenicurus* |  | *Phalangiidae* |
|  | *Pica pica* |  | *Trombidiidae* |
|  | *Phylloscopus collybita* |  | *Thomisidae* |
|  | *Phylloscopus trochilus* |  | *Staphylinidae* |
|  | *Sitta europaea* |  |  |
|  | *Turdus merula* |  |  |
| Sum | 61 | 162 | 652 |
| **Paloheinä (Helsinki)** | *Cyanistes caeruleus* | *Acer sp.* | *Araneidae* |
|  | *Columba palumbus* | *Alnus incana* | *Carabidae* |
|  | *Dendrocopos major* | *Betula pendula* | *Formicidae* |
|  | *Erithacus rubecula* | *Betula pubescens* | *Linyphiidae* |
|  | *Fringilla coelebs* | *Picea abies* | *Lycosidae* |
|  | *Parus major* | *Pinus sylvestris* | *Nemastomatidae* |
|  | *Phylloscopus sibilatrix* | *Populus tremula* | *Phalangiidae* |
|  | *Phylloscopus trochilus* | *Prunus padus* | *Trombidiidae* |
|  | *Phylloscopus trochiloides* | *Salix aurita* | *Tipuloidea* |
|  | *Regulus regulus* | *Sorbus aucuparia* | *Thomisidae* |
|  | *Sylvia atricapilla* |  | *Staphylinidae* |
|  | *Troglodytes troglodytes* |  |  |
|  | *Turdus merula* |  |  |
|  | *Turdus philomelos* |  |  |
|  | *Turdus viscivorus* |  |  |
| Sum | 136 | 73 | 460 |
| **Viikki**  **(Helsinki)** | *Alauda arvensis* | *Acer sp.* | *Araneidae* |
|  | *Anthus trivialis* | *Betula pendula* | *Carabidae* |
|  | *Corvus corax* | *Betula pubescens* | *Coccinellidae* |
|  | *Corvus cornix* | *Picea abies* | *Elateridae* |
|  | *Cyanistes caeruleus* | *Pinus sylvestris* | *Formicidae* |
|  | *Columba palumbus* | *Populus tremula* | *Gnaphosidae* |
|  | *Dendrocopos major* | *Prunus padus* | *Linyphiidae* |
|  | *Emberiza citrinella* | *Sorbus aucuparia* | *Lycosidae* |
|  | *Erithacus rubecula* | *Tilia cordata* | *Nemastomatidae* |
|  | *Hippolais icterina* |  | *Phalangiidae* |
|  | *Motacilla alba* |  | *Trombidiidae* |
|  | *Muscicapa striata* |  | *Tipuloidea* |
|  | *Motacilla flava* |  | *Thomisidae* |
|  | *Parus major* |  | *Staphylinidae* |
|  | *Pica pica* |  |  |
|  | *Phylloscopus collybita* |  |  |
|  | *Passer domesticus* |  |  |
|  | *Phylloscopus sibilatrix* |  |  |
|  | *Phylloscopus trochilus* |  |  |
|  | *Sylvia atricapilla* |  |  |
|  | *Troglodytes troglodytes* |  |  |
|  | *Turdus merula* |  |  |
|  | *Turdus philomelos* |  |  |
|  | *Turdus pilaris* |  |  |
| Sum | 100 | 67 | 554 |
| **Total abundance** | **695** | **586** | **2510** |

**Table S2**. Environmental data and how they were measured.

| **Variable name & unit** | **How it was measured** | **Range of values per plot** |
| --- | --- | --- |
| Number of individual trees | We counted the individual number of tree stands, both alive and dead, found within the 5 m radius of the measuring point.  We only included trees that had stems >10 cm in diameter. Trees with stems <10 cm diameter were included in the ground vegetation cover estimates.  If the roots of the species were based outside the radius, we did not include it in our survey. | 0-18 trees |
| Number of tree species | We counted the number of tree and bush species found within the 5 m radius of the measuring point. If the tree was dead but could still be recognized to its species, we included it in our species count.  Vegetation outside the radius was not considered. If the roots of the species were based outside the radius, we did not include it in our survey. | 2-9 species |
| Tree stand height (m) | We measured the height of tree stands within the measuring point using a relascope.  We only included trees that had stems > 10cm in diameter. Trees with stems < 10cm diameter were included in the ground vegetation cover estimates.  If the roots of the species were based outside the radius, we did not include it in our survey. | 0-36m |
| Tree stand diameter (cm) | We measured the diameter of tree stands within the measuring points radius at chest height (~130cm).  We only included trees that had stems > 10cm in diameter. Trees with stems < 10cm diameter were included in the ground vegetation cover estimates.  If the roots of the species were based outside the radius, we did not include it in our survey. | 0-90cm |
| Ground vegetation cover (%) | We estimated the percentage of trees with stem diameters < 10cm that covered the 5m radius inside the measuring point.  Bushes and other vegetation at 30cm height or above were included. Grass and flowers < 30cm height was excluded. | 0%-100% |
| Basal area cover of tree stands (m^2^/ha) | Standing at the centre of the measuring plot, we used a relascope (1m fibreglass stick with 2cm wide internal cross piece) and counted all trees in a 360° survey whose stems/trunks fitted within the cross piece.  The dimensions of the relascope gives a ratio 1:50 and basal area factor of 1, meaning every counted tree represents 1 m^2^/ha  If the tree stem barely filled the cross piece, every other type of this stem/trunk was counted. If the stem/trunk couldn’t fill the cross piece it was excluded from the count. | 3m^2^/ha – 42m^2^/ha |
| Proportion of dead trees (%) | We divided the number of dead trees (standing and/or logs) with the number of alive trees found within the measuring points radius to get a proportion of deadwood. | 0%-100% |

**Table S3.** Guidelines of the 11^th^ Finnish National Forest Inventory (Valtakunnan Metsien Inventointi, VMI11 (2009)) and the decaying wood project of Tapio Oy (2023) used for assessing the natural condition of forests and stands*.*

1. Forest structure (VMI11, 2009)

1 = Natural or similar to natural. The spatial distribution of the forest is random and the size of the forest varies. The canopy is layered and trees from the previous generation are present here and there. The pattern may show minor signs of old harvesting, but they have not affected the spatial distribution, structure or tree species composition of the forest. The dominant forest is old, at least of silvicultural regeneration age. After complete storm damage, an opening or a naturally regenerated sapling can belong to this category, if the damage has affected a stand that is ripe for regeneration and the trees that died as a result of the damage have not been removed. The category also includes natural areas of recent fire and young forests that have developed naturally after a forest fire, as well as new, natural landforms that have emerged from the sea as a result of land uplift.

2 = The stand has emerged naturally, the structure deviates slightly from the natural state or there are signs of minor thinning or removal of overhanging trees.

3 = The stand is uniform in spatial arrangement and one-sided in tree species and size distribution, e.g. as a result of cultivation or thinning.

2. Deadwood continuum (Tapio Oy, 2023)

The amount of dead standing and groundwood is estimated in relation to the potential amount of deadwood in the natural forest of the growth site in question.

0 = No deadwood

1 = Weak deadwood continuity; the majority of the dead trees in the figure are either dead standing or hard groundwood, with older groundwood trees present only once.

2 = Moderate deadwood continuity; in addition to dead standing and hard groundwood, at least about a third of the dead trees in the figure are more decayed (medium decayed and deeply decayed) groundwood.

3 = Good deadwood continuity; in the figure, at least about a third of the groundwood trunks are deeply decayed, at least about a third are moderately decayed and at most about a third are still hard, fairly recently dead.

**S3. Analysis**

**Table S4.** Summary of models, their ecological effects examined, their responses, fixed factors, random factors and families. * signifies that both the main effects of variables and their interactions were considered. + signifies that only main effects where considered.

| **Model** | **Response variable** | **Fixed factors** | **Random factors** | **Family** |
| --- | --- | --- | --- | --- |
| 1 | Initial survival per prey item | Group size * Colouration * Palatablity | Year \| Transect | Binomial |
| 2 | Initial survival | Number of invertebrate predators + Number of invertebrate predator families | Year \| Transect | Binomial |
| 3 | Initial survival | Basal area cover + Ground vegetation cover + Number of tree species + Tree height + Deadwood percentage | Year \| Transect | Binomial |
| 4 | Initial survival | Forest structure + Deadwood continuum | Year \| Transect | Binomial |
| 5 | Number of invertebrate predators | Forest structure +  Deadwood continuum | Year \| Transect | Negative-binomial (nbinom1) |
| 6 | Number of invertebrate predator families | Forest structure +  Deadwood continuum | Year \| Transect | Gaussian |
| 7 | Initial survival | (Group size + Colouration + Palatability) * (Number of invertebrate predators + Number of invertebrate predator families) | Year \| Transect | Binomial |
| 8 | Initial survival | (Group size + Colouration + Palatability) * (Basal area cover + Ground vegetation cover + Number of tree species + Tree height + Deadwood percentage) | Year \| Transect | Binomial |
| 9 | Initial survival | (Group size + Colouration + Palatability) * (Forest structure + Deadwood continuum) | Year \| Transect | Binomial |

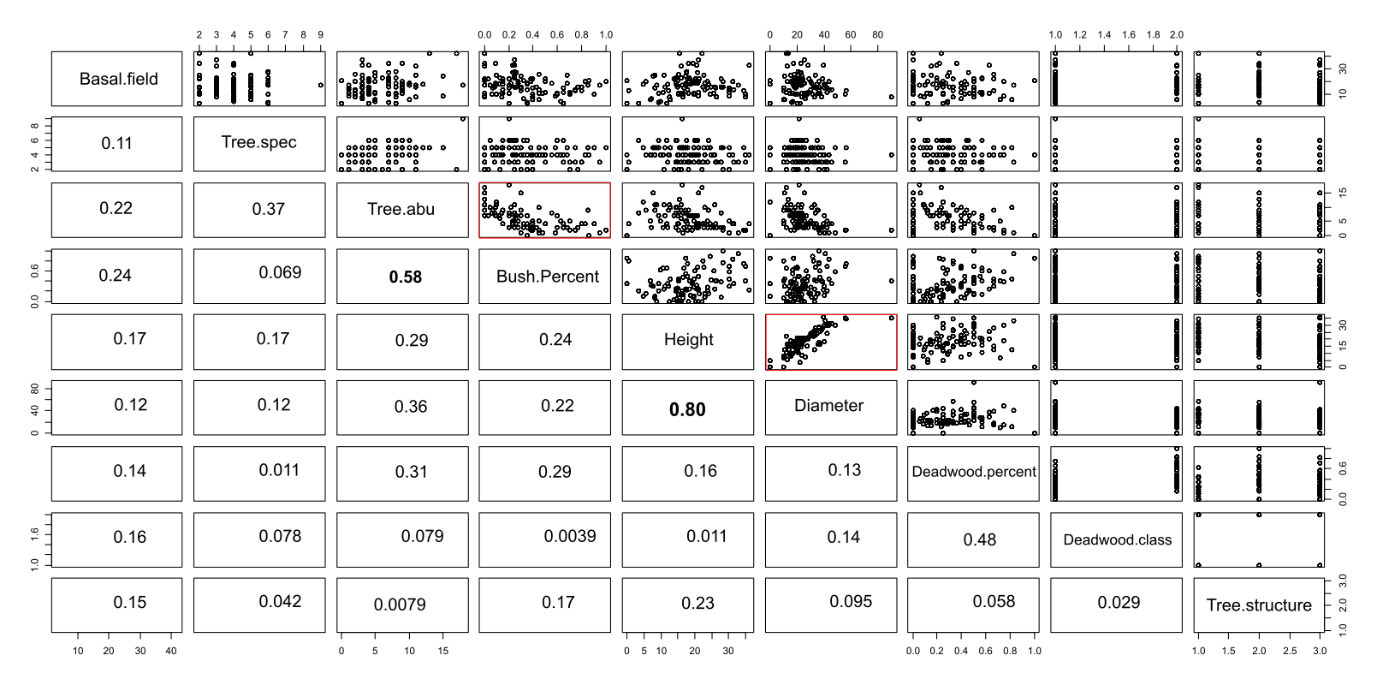

**Fig S8** Correlation table of environmental variables. All variables showed significant correlations with each other from using the cor.test() function in R. When examining figures visually reveals that most correlations do not appear as meaningful. We excluded variables that exceeded 0.5 (marked bold and highlighted in red rectangles) in their correlation coefficients as these appeared to show correlations when visually examined.
